# Toward a model of uORF-mediated translational control: An integrated bioinformatic and experimental approach

**DOI:** 10.1101/2025.07.28.667205

**Authors:** Britt Hanson, Nenad Svrzikapa, Ning Feng, Hans J. Friedrichsen, Angus J. Lennaárd, Rushdie Abuhamdah, Nicole Hemmer, Katarzyna Chwalenia, Matthew Drake, Yahya Jad, Samir EL Andaloussi, Matthew J.A. Wood, Thomas C. Roberts

## Abstract

Upstream open reading frames (uORFs) are short translated regions that occur in the 5□ untranslated regions (5□ UTRs) of mRNA transcripts where they primarily serve to repress expression translation of the downstream primary open reading frame (pORF). Their widespread presence across mammalian transcriptomes suggests an important role in shaping the proteome, although the mechanistic basis of their regulatory effects remain incompletely understood. Here we present an integrated experimental and computational investigation into the features that govern uORF-mediated translation control. Using high-resolution proteomics data from 29 healthy human tissues and machine learning-based simulations, we have systematically dissected how features including uORF length, amino acid composition, start codon position, stop codon position, and Kozak context influence repressive activity, and performed experimental validation using reporter gene constructs. We also investigated how multiple uORFs within a single 5□ UTR can interact in synergistic or antagonistic ways, with the potential to produce counterintuitive effects on pORF translation. From these studies, we present a model of uORF function, suggesting a hierarchy of uORF feature importance, and proposing that a combination of uORF translation initiation probability, ribosome recycling rate, intercistronic ternary complex recharging requirements, and ribosome stalling mechanisms underlie uORF repressive activity. Together, these studies provide a comprehensive view of the molecular logic underlying uORF activity, offering new insights into their endogenous and highlighting their potential as targets for drug development.

## Introduction

In eukaryotic mRNAs, the primary open reading frame (pORF) is often preceded by one or more upstream open reading frames (uORFs). A uORF consists of an initiation codon (typically an AUG) within the 5□ untranslated region (5□ UTR) of protein-coding mRNA transcripts, followed by an in-frame stop codon. uORFs are highly diverse in terms of sequence length, number of uORFs per transcript, distance from the 5□ m^7^G-cap, distance from the pORF, strength of Kozak context sequence, evolutionary conservation, and whether the uORF overlaps with the pORF. uORFs act to regulate gene expression in several distinct ways. The predominant mechanism appears to be the regulation of translation, whereby the presence of a uORF results in a reduction in the translation of the downstream pORF.^1–5^ However, other mechanisms have also been reported including nonsense mediated decay,^6^ and altering isoform usage.^7^ As such, inhibition of uORF translation may result in activation of pORF expression in many cases. The widespread occurrence of uORFs suggests that a large proportion of the transcriptome may be held in a semi-repressed state, the function of which is possibly to control the translation of dosage-sensitive genes and thereby prevent toxic or wasteful protein overproduction. Furthermore, uORFs are to some extent subject to dynamic regulation, as stress-induced phosphorylation of eIF2α (eukaryotic initiation factor-2) can increase leaky scanning by the 40S ribosome subunit, leading to stimulus-dependent relief of uORF-mediated repression.^8–10^ Additionally, uORFs are translated into peptides,^11^ which may themselves exhibit biological functions.^12–16^ Overall, uORFs constitute an important component of the gene regulatory apparatus of the cell.

Genetic variants that introduce or disrupt a uORF start or termination codon, or modify the strength of the uORF Kozak sequence context, can have important consequences for human disease.^1,17^ For example, a common variant (rs1801020) in coagulation factor XII (*F12*), present in 20% of Caucasian and 70% of Asian populations, disrupts a uORF proximal to the *FXII* translation initiation site (TIS) resulting in a predisposition to thrombosis.^1,18–22^ uORFs are predicted to exist in a large proportion of proto-oncogenes,^17,23–25^ with loss-of-function uORF mutations identified in several human malignancies.^26^ Disease-relevant uORFs have also been identified in genes associated with Alzheimer’s disease (i.e. *BACE1*),^27^ asthma,^24^ type 2 diabetes (*KCNJ11*, *ARAP1*),^24^ and Parkinson’s disease (*PARK7*).^24^

Molecular medicine approaches that can disrupt the function of specific uORFs may be beneficial in the treatment of autosomal dominant haploinsufficiency conditions, or for the translational upregulation of disease modifying genes. As such, the targeting of uORF sequences with steric block antisense oligonucleotides (ASOs) was reported to augment the translation of both human (*RNASEH1*, *SFXN3*) and mouse (*Mrpl11*, *Lrpprc*) genes.^28^ However, efforts from our group to reproduce these findings for *RNASEH1* have been unsuccessful.^29^ Nevertheless, other approaches for uORF interference have been reported, including ASO-mediated disruption of RNA secondary structure-uORF interactions,^30^ exon skipping to exclude uORF containing exons^31^, and CRISPR-mediated editing of uORF start codons via indel formation^32–34^ or base editing.^35^

Given the emerging importance of uORFs, we sought to systematically investigate how the properties of uORFs influence their repressive activity. Previous studies have investigated how uORF features influence repressive activity using various approaches.^1,36–43^ Here we describe an integrated approach to investigating uORF features using a combination of computation analyses (using high-resolution proteomics data from 29 healthy human tissues^44^ and machine learning-based simulations^45^) and experimental validation using reporter gene constructs (primarily using *HOXA11* as an exemplar). This comprehensive strategy was applied to various uORF features including length, amino acid composition, Kozak sequence context, start codon position, stop codon position, uORF-pORF intercistronic distance, and the effects of multiple uORFs in series, which has revealed new insights into uORF activity. Building on the resulting findings, we present a working model of human uORF function.

## Results

### Prediction of uORFs in the human and mouse transcriptomes

The 5□ UTRs of all human (hg38) and mouse (mm10) protein-coding RefSeq transcripts were interrogated using a custom script to predict uORF occurrences (**Figure S1**). A total of 25,503 (59.3%) human, and 14,479 (48.5%) mouse transcripts were found to have at least one predicted uORF (**Figure 1A**), which was comparable to previous estimates using earlier genome builds.^1,46–48^

**Figure 1.**
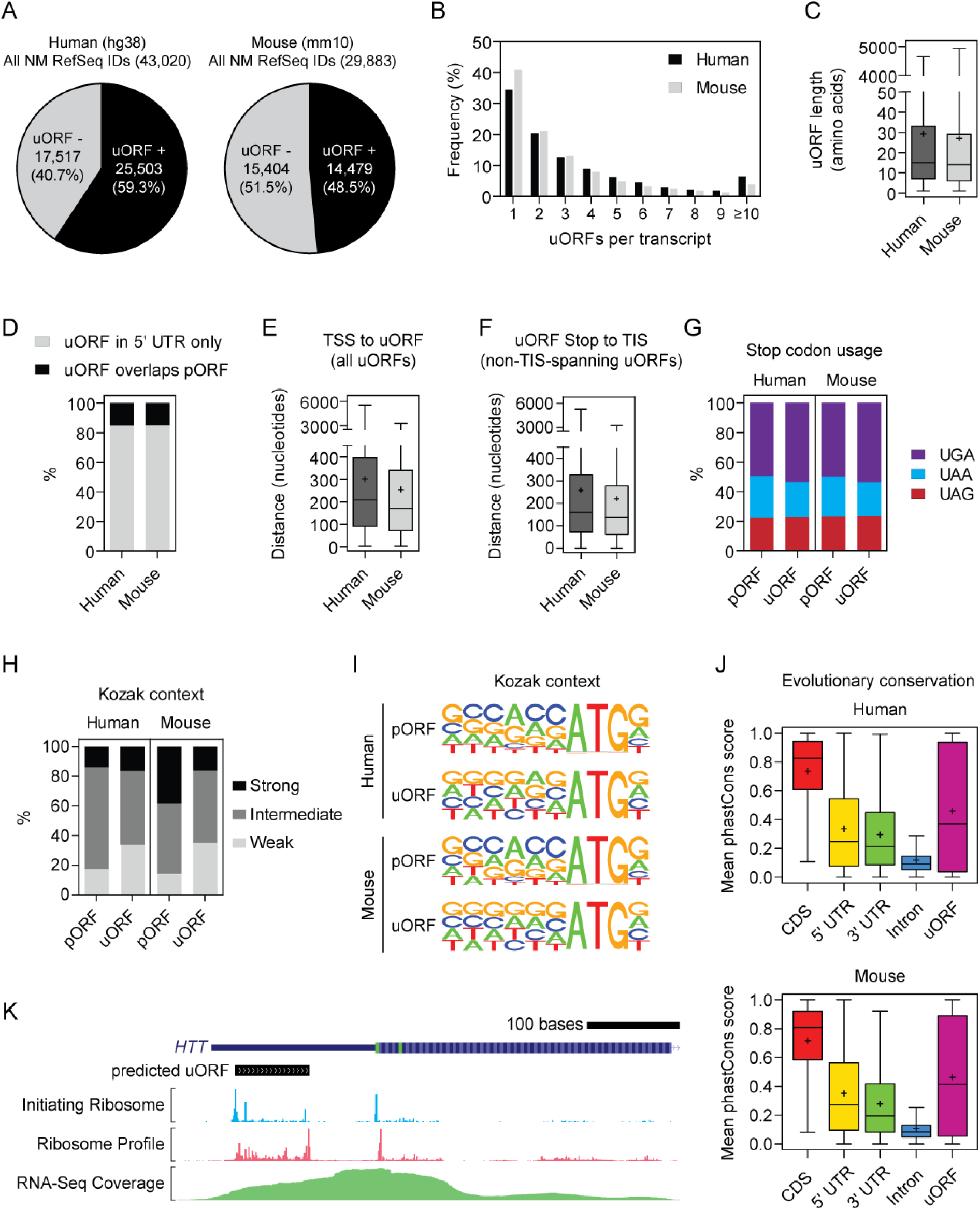
Prediction and characterization of human and mouse uORFs. Human (hg38) and mouse (mm10) transcripts were searched for potential uORFs with ATG start codons. (**A**) Pie charts illustrating the proportion of transcripts containing at least one predicted uORF. (**B**) Distribution of the number of predicted uORFs per transcript. (**C**) Length distributions for predicted uORF peptides. (**D**) Proportion of uORFs which overlap with the pORF. (**E**) Box plots showing the distances between (**E**) the Transcription Start Site (TSS) and the predicted uORF, and (**F**) the end of the predicted uORF and the start of the pORF (for non-pORF-spanning uORFs only). (**G**) Stop codon usage for pORFs and uORF predictions. The strength of the Kozak sequence context surrounding the start codon for pORFs and predicted uORFs was (**H**) scored as ‘Strong’, ‘Intermediate’, or ‘Weak’ and (**I**) visualized as sequence logos. (**J**) PhastCons scores for uORF prediction relative to other transcript features. (**K**) Visualization of a single uORF at the *HTT* locus together with aggregate initiating ribosome profile, ribosome footprint, and RNA-Seq coverage. Box plots show the median and interquartile range, whiskers represent the minimum and maximum values, and the mean value is represented by ‘+’.

Transcripts with predicted uORFs were categorized depending on the number of uORFs per transcript, with single uORF transcripts constituting the largest group (∼34% human, ∼41% mouse), meaning that the majority of uORF-containing transcripts harbor more than one uORF (**Figure 1B**). Indeed, ∼6% of human and ∼4% of mouse transcripts contained 10 or more predicted uORFs (**Figure 1B**). The majority of predicted uORFs were short, with an interquartile range of ∼6 to ∼33 amino acids in length (**Figure 1C**), and ∼85% of uORFs did not overlap with the pORF (**Figure 1D**). Distances between the transcription start site (TSS) and the start of the uORF, and between the uORF stop codon and the translation initiation sites (TIS, i.e. the pORF) were similar for both human and mouse (**Figure 1E,F**). The most utilized stop codon was UGA for both pORFs and uORFs for the human and mouse transcriptomes (**Figure 1G**).

The Kozak context sequence was generally weaker for uORFs than for pORFs (**Figure 1H,I**). A ‘strong’ Kozak consensus sequence is defined as having both a G at the +4 position and a purine at the −3 position.^2^ Genomic coordinates were calculated for predicted uORFs which enabled average phastCons scores^49^ to be determined for each uORF (and compared with 5□ UTR, 3□ UTR, CDS, and intronic regions). Conservation values for uORF sequences were highly diverse, essentially spanning the full range of possible phastCons values, suggesting that some uORFs are highly conserved, while others are poorly conserved (**Figure 1J**). Overall, predicted uORF features were highly similar between human and mouse (**Figure 1A-J**). uORF predictions were visualized using the GWIPS-viz genome browser^50^ together with aggregated Ribo-Seq and RNA-Seq data, which allowed for the identification of uORFs for which there is experimental evidence. The *HTT* gene is shown as an example (**Figure 1K**) whereby a prominent initiating ribosome peak and ribosome footprint are observed at the predicted uORF.

### Validation of predicted uORF repressive activity

In order to select candidates for validation, predicted human uORF-containing genes were intersected with the COSMIC Cancer Gene Census list of oncogenes and tumor suppressors^51,52^ and the list further restricted by filtering out genes containing more than one uORF, or where the uORF overlaps with the pORF (**Figure S2**). Candidate uORF-containing genes (*JUN*, *IKBKB*, *KDR*, *MAP2K2*, *SMO*, *KDM5A*, *NCOR1*, *FOXL2*, *MAP3K1*, *HOXA11*, and *HOXA9,* **Figure 2A**) were tested by cloning the corresponding 5□ UTR upstream of *Renilla* luciferase (RLuc) in a dual luciferase reporter system (**Figure S3A**). The 5□ UTRs of *KDR*, *SRY*, and *RNASEH1* were selected based on previous reports.^1,28,29^ For each candidate gene, control constructs were generated in which the uORF was disrupted by mutagenesis of the upstream ATG (uATG) to TTG. A significant increase in RLuc expression was observed as a result of uORF disruption for 9 of 13 of the 5□ UTRs tested from this panel (**Figure 2B**). The *SMO* 5□ UTR exhibited the highest level of uORF-mediated translational repression, whereby inactivating its uORF resulted in an ∼11-fold increase in reporter gene activity relative to the wild-type (WT) control (*P*<0.001). RLuc expression was increased by 4.4-fold (*P*<0.001) following uORF disruption of the *HOXA11* 5□ UTR, and between 1.3 and 3.8-fold for the remaining significantly de-repressed 5□ UTRs. For the other four candidate 5□ UTRs, uORF disruption resulted in an increase in RLuc expression (between 1.2 and 2-fold) that did not reach significance at the *P*<0.05 level. Luciferase transcript levels were largely unchanged between mutant and WT constructs, indicating that uORF-mediated regulatory effects manifest at the level of translation in these constructs (**Figure 2C**).

**Figure 2.**
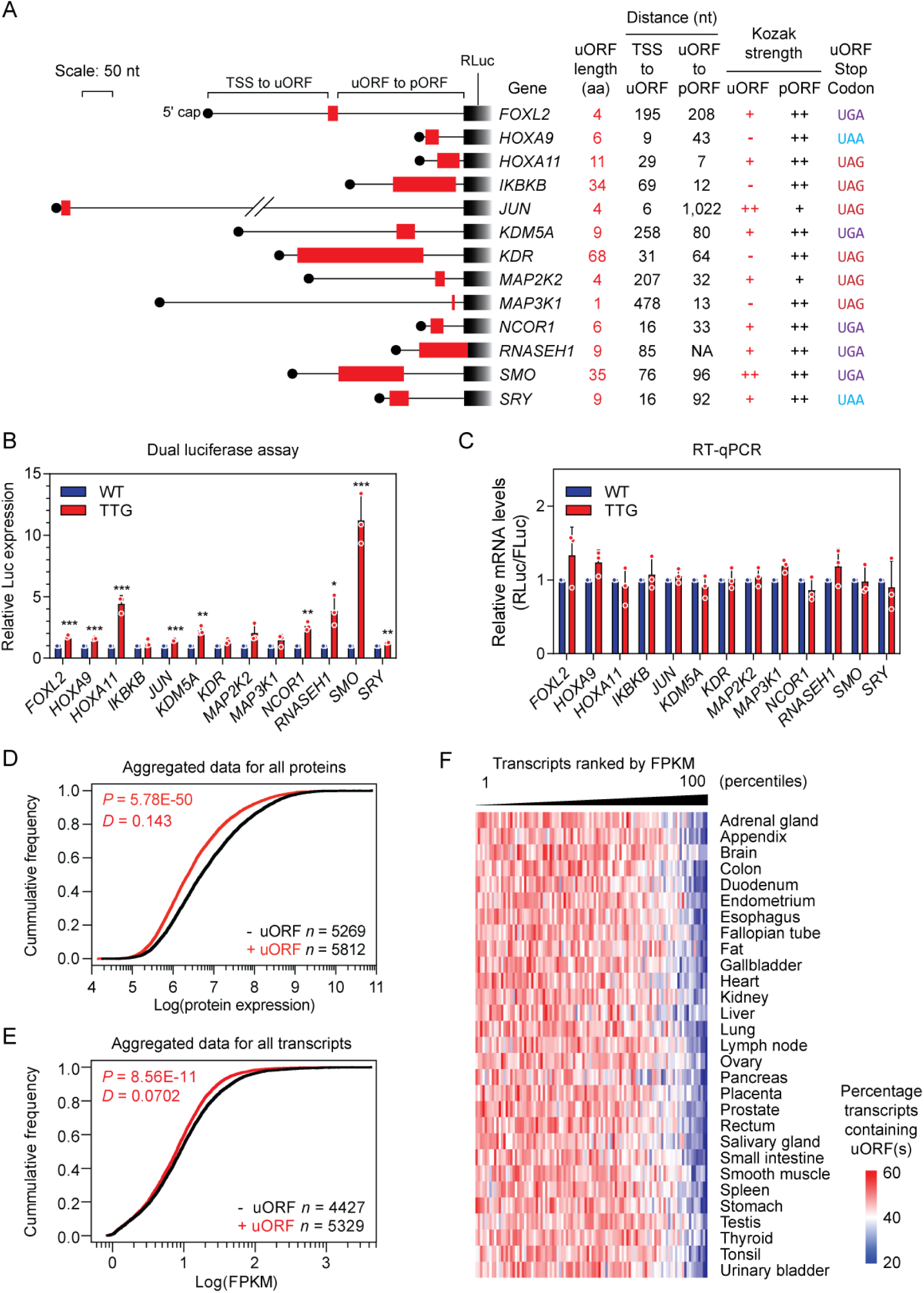
Experimental validation of uORF-mediated translational repression. (**A**) Features of the 5□ UTRs of a panel of oncogenes and tumor suppressor genes which were cloned upstream of RLuc in a dual-luciferase reporter (DLR) vector. Each 5□ UTR contains a single predicted uORF. uORFs are indicated by red boxes. uORF-containing wild-type (WT) and uORF-knockout (TTG, in which the uORF ATG was mutated to TTG) versions of each pDLR vector were transfected into HEK293T cells (*n*=3 independent experiments) and (**B**) a DLR assay performed to assess the effect of uORF activity on *Renilla* luciferase (RLuc) protein expression. (**C**) RT-qPCR of RLuc mRNA transcript levels, normalized to FLuc. Values of the WT and TTG version of each uORF were scaled to a value of 1, and are mean+SD. Statistical significance was determined by a Student’s *t*-test relative to the corresponding gene-specific WT control, \**P*<0.05, \*\**P*<0.01, and \*\*\**P*<0.001. Publicly available matched proteomics and RNA-Seq data from 29 healthy human tissues^44^ were analysed and cumulative distributions plotted for (**D**) protein, and (**E**) RNA expression for transcripts containing one or more predicted uORFs versus transcripts containing no uORFs. Statistical significance between distributions was tested by Kolmogorov-Smirnov test. (**F**) Heatmap of predicted uORF frequency across mRNA expression bins in all 29 human tissues. Highly abundant transcripts tend to lack uORFs.

We also investigated uORF-containing genes that are associated with neuromuscular disorders, and might constitute therapeutic targets. To this end, we identified single uORFs in the *SMN2* and *UTRN* genes, upregulation of which is expected to be beneficial in the cases of spinal muscular atrophy and Duchenne muscular dystrophy, respectively.^53,54^ *SMN2* and *UTRN* WT 5□ UTR and TTG mutant constructs were generated and tested for luciferase activity. Interestingly, both of these uORFs were located in close proximity to the 5□ cap, and neither uORF was found to exert a repressive effect on its respective pORF (**Figure S4**).

To assess the impact of uORF sequences on translational repression on a global scale, we utilized publicly available, matched proteomics/transcriptomics data from 29 healthy human tissues (for 18,072 transcripts and 13,640 proteins).^44^ Data were filtered to exclude genes with multiple transcript isoforms where at least one of those isoforms lacked a uORF, and the remaining data binned as ‘uORF-containing’ or ‘uORF-lacking’ respectively. The cumulative distribution function (CDF) of protein expression data (aggregated across all tissues) for all genes in each bin were plotted and the difference between the distributions tested by Kolmogorov-Smirnov test. Consistent with previous reports, we observed that genes containing uORFs (*n*=5,269) have significantly lower median protein expression levels than genes lacking uORFs (*n*=5,812) (*P*=5.78×10^−50^, *D*=0.143, **Figure 2D**). Separate CDF analyses of each individual tissue yielded very similar results for both protein and RNA expression (**Figure S5**). An equivalent CDF analysis was performed using aggregated transcript expression data, which showed that uORF-containing transcripts (*n*=5,329) tended to be expressed at lower levels than transcripts without uORFs (*n*=4,427), although the effect size was smaller than that observed for protein expression (*P*=8.56×10^−11^, *D*=0.0702) (**Figure 2E**). Notably, the top 10% most highly expressed transcripts (highest fragments per kilobase of exon per million mapped fragments, FPKMs) had a tendency to lack uORFs, which was the case for all 29 tissues (**Figure 2F**). Taken together, these data suggest that the presence of one or more uORFs is associated with a reduction in protein expression, and to a lesser extent, a reduction in transcript expression.

### Analysis of uORF structure and function

The *HOXA11* uORF was selected for further study, as this uORF conferred a strong translational repressive effect in reporter studies (**Figure 2B**), both the 5□ UTR and uORF are of convenient lengths for facile experimental manipulation, and this uORF was also previously demonstrated to be function in a murine context.^55^ Furthermore, publicly available Ribo-Seq data revealed a prominent initiating ribosome peak at the uORF AUG codon and a high ribosome footprint read density at the predicted uORF (**Figure 3A**). The activity of the *HOXA11* uORF was confirmed in a cell-free *in vitro* translation model system using synthetic mRNA (**Figure 3B**).

**Figure 3.**
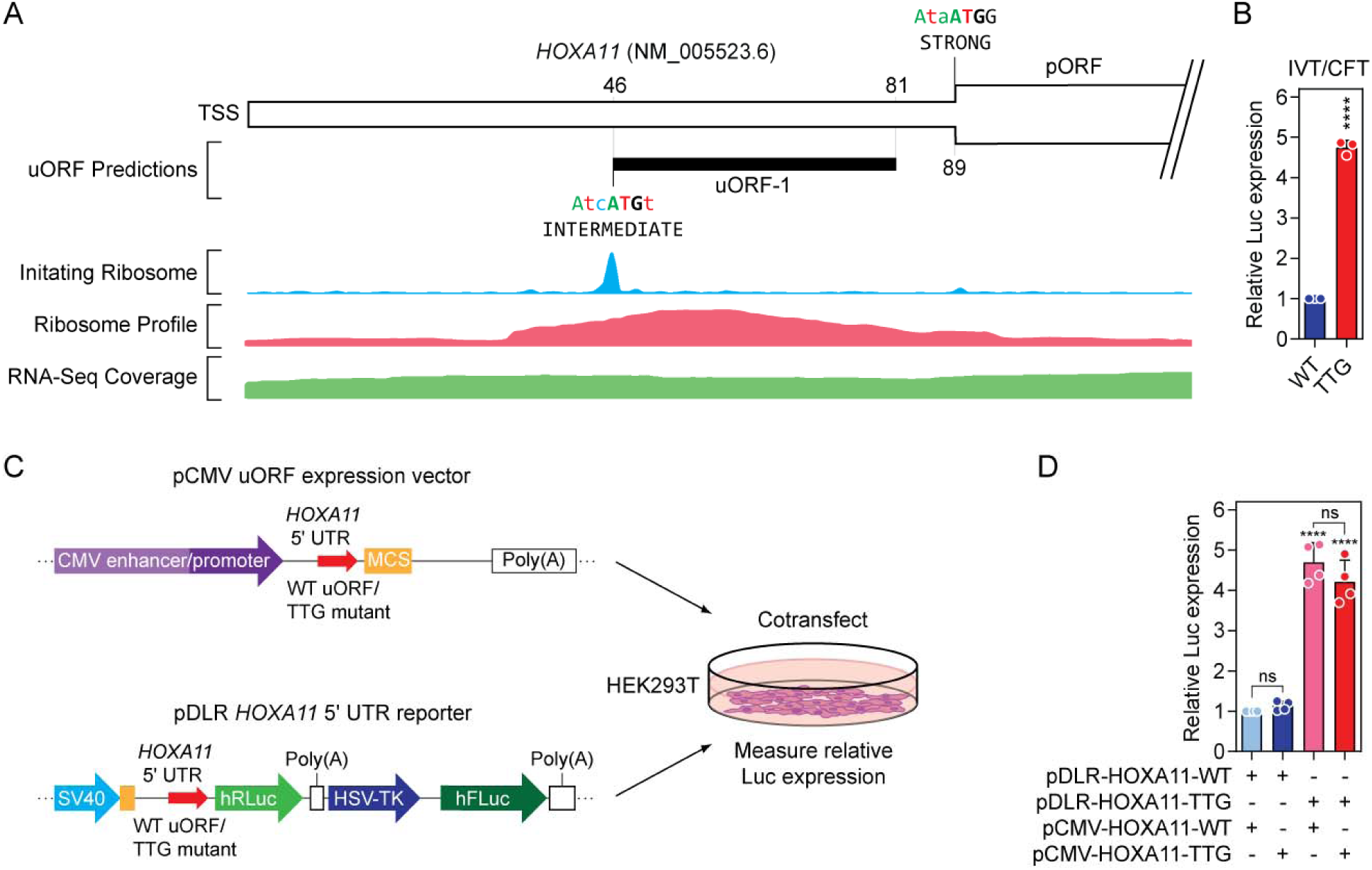
*HOXA11* is an archetypal uORF regulated gene. (**A**) Schematic of the *HOXA11* uORF with overlaid aggregated Ribo-Seq and RNA-Seq data. (**B**) uORF activity was assessed by *in vitro* transcription/cell-free translation (IVT/CFT) assay using synthetic mRNA transcripts encoding the Renilla luciferase transgene following either the WT *HOXA11* 5□ UTR (WT) or a mutant construct in which the *HOXA11* uORF was disrupted by mutating its start codon (TTG). (**C**) A plasmid encoding the HOXA11 uORF peptide sequences driven by the cytomegalovirus (CMV) promoter (or non-translated TTG control) was co-transfected into HEK293T cells together with dual luciferase reporter plasmids with either the WT *HOXA11* 5□ UTR or the uORF disrupted TTG control. (**D**) Luciferase activity was determined 24 hours post transfection, whereby no uORF peptide-mediate translational repression effects were observed *in trans*. Values are mean+SD, (*n*=3-4 independent experiments), and were scaled such that the mean of the control group was returned to a value of 1. Statistical significance was assessed by Student’s *t*-test or one-way ANOVA with Bonferroni *post h*oc test, as appropriate. \*\*\*\**P*<0.0001, ns=not significant.

Previous work has identified functional roles for uORF-encoded micropeptides in certain cases.^12–16^ To investigate this possibility, constructs were generated in which the *HOXA11* uORF was tagged at the C-terminus with either a single FLAG tag or with a HiBiT tag. Tagged *HOXA11* uORF peptides were undetectable by either anti-FLAG western blot or HiBiT biocomplementation assay (data not shown). To test whether the *HOXA11* uORF sequence can mediate its repressive effects in *trans*, the DNA encoding the *HOXA11* uORF peptide (or the TTG mutant) was cloned into an RNA Polymerase II expression vector, and co-transfected with the *HOXA11* 5□ UTR TTG mutant dual luciferase reporter (or WT construct) (**Figure 3C**). No reporter downregulation activity was observed using this assay (**Figure 3D**). Taken together, these findings suggest that the *HOXA11* uORF is a *cis* regulator of downstream pORF expression, and that its encoded micropeptide is likely unstable, and rapidly degraded after synthesis. We therefore used *HOXA11* as a model system for studying uORF function as an archetypal example of uORF-mediated translational repression.

We next investigated the contributions of fundamental uORF properties on translation repression activity. To this end, a plasmid construct was generated in which the *HOXA11* uORF was replaced with a Golden Gate cloning site (**Figure S3B**), and a variety of mutants of the WT *HOXA11* uORF were generated.

The native *HOXA11* uORF Kozak consensus is weak. Altering the Kozak consensus to increase its strength resulted in an enhancement of downstream gene repression by ∼17% which did not reach statistical significance at the *P*<0.05 level (**Figure 4A**). Consistently, global analysis of publicly available protein expression data revealed a small shift towards greater repression (*D*=0.063) in transcripts containing Strong (*n*=341) versus transcripts containing Weak (*n*=1,705) Kozak uORFs which did not reach statistical significance (**Figure 4B**).

**Figure 4.**
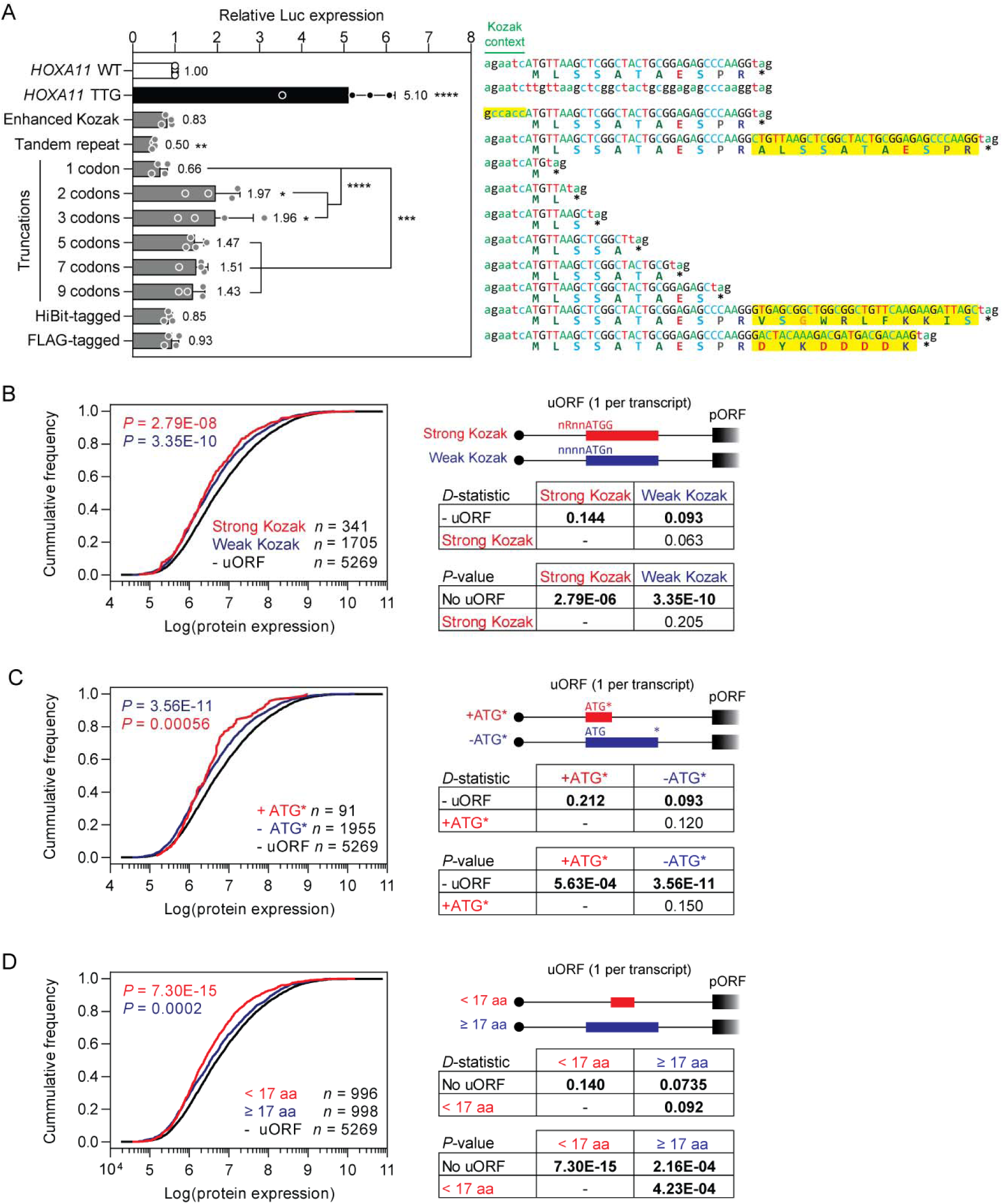
Effect of uORF length and Kozak context sequence on uORF activity. (**A**) The *HOXA11* 5□ UTR dual luciferase reporter was mutated to enhance the uORF Kozak consensus sequence, to alter the length of the uORF, or to tag the uORF peptide with FLAG or HiBiT tags. These mutant constructs were transfected in HEK293T cells and luciferase activity measured 24 hours later. Key mutated bases are highlighted in yellow. Aggregated proteomics data from 29 healthy human tissues were binned according to (**B**) the strength of the uORF Kozak consensus sequence, (**C**) minimal uORF (ATG-Stop) versus other uORFs, and (**D**) uORF length (<17 amino acids versus ≥17 amino acids). In each case, data are presented as cumulative distribution function plots. Statistical differences between distributions were assessed by Kolmogorov-Smirnov test and *D*-statistics and *P*-values tabulated. The numbers of proteins in each bin are indicated. Values are mean+SD, (*n*=4 independent experiments), and were scaled such that the mean of the WT control group was returned to a value of 1. Statistical significance was assessed by one-way ANOVA and Bonferroni *post hoc* test. \**P*<0.05, \*\**P*<0.01, \*\*\**P*<0.001, \*\*\*\**P*<0.0001.

We subsequently investigated the importance of uORF peptide length through a series of truncations from the C-terminus (i.e. to generate uORFs that were 1, 2, 3, 5, 7, and 9 amino acids in length). Progressive truncation of the *HOXA11* uORF resulted in a loss of repressive activity that was proportional to the degree of truncation (**Figure 4A**). Notably, the 2 and 3 codon uORF variants exhibited ∼2-fold de-repression relative to the WT *HOXA11* uORF (*P*<0.05). However, the minimal uORF (i.e. 1 codon, ATG-STOP, M*, methionine-STOP) did not fit this pattern and instead exhibited repression that was not statistically different from the WT *HOXA11* uORF but was ∼3-fold reduced relative to the 2-codon construct (*P*<0.0001) (**Figure 4A**). It has been reported that an unoccupied ribosome exit (E) site is associated with translational arrest,^56^ which might account for the repressive effects of the minimal uORF. Such minimal uORFs occur frequently in our prediction datasets; 5,447 (6% of all uORFs) in human, and 2,759 (6.5%) in mouse, although often in multi-uORF contexts. Notably, these uORFs have been overlooked in previous studies based on arbitrary uORF definitions (e.g. that they consist of at least nine nucleotides).^1^ Accordingly, global analysis of aggregated human proteomics data revealed that of all single uORF-containing transcripts, those which contained the minimal uORF (*n*=91) tended to be expressed at lower levels than those that contained other types of uORFs (*n*=1,955), although this effect did not pass the threshold for statistical significance (*D*=0.12, **Figure 4C**).

Increasing the length of the uORF by repeating the entire amino acid sequence resulted in a significant 2-fold (*P*<0.01) increase in repressive activity. By contrast, extending the length of the uORF by addition of tag sequences (i.e. FLAG and HiBiT) had only minimal effects on the degree of translational repression (**Figure 4A**). Aggregated human proteomics data were interrogated to determine the global effect of uORF length. Single uORF transcripts were binned according to whether they contained ‘Short’ (i.e. <17 amino acids, *n*=996) or ‘Long’ (i.e. ≥17 amino acids, *n*=998) uORFs and CDFs plotted. This binning strategy was selected in order to maximize the number of transcripts per bin (and therefore also statistical power). Short uORFs were found to be more repressive than Long uORFs (**Figure 4D**) (*D*=0.092, *P*=4.23×10^−4^).

Changes in luciferase expression in these various reporter constructs (**Figure 4A**) could not be explained by alterations in transcript levels (**Figure S6**).

### Effect of uORF reading frame on repressive activity

The relative reading frames for the *HOXA11* uORF and pORF are offset by 1 base pair. Mutant constructs were generated whereby additional non-coding ‘stuffer’ bases were added such that this offset was altered to 2 base pairs, and 0 base pairs (i.e. in the same reading frame as the pORF). Changing the relative *HOXA11* uORF reading frame did not alter uORF repressive activity (**Figure S7A**). Global analysis of relative uORF/pORF reading frames revealed that the uORFs in frame 0 tended to be the least repressive relative to transcripts without uORFs (*D*=0.0836, *P*=0.00273, *n*=513), in contrast with uORFs in frame 1 (*D*=0.121, *P*=1.7×10^−8^, *n*=709) and frame 2 (*D*=0.131, *P*=6.35×10^−10^, *n*=717) (**Figure S7B**). uATGs that are in frame with the pORF have the potential to generate N-terminally extended versions of the pORF-encoded protein, which likely accounts for their relatively lower level of repressive activity. Notably, upstream ATGs that share a termination codon with the pORF were computationally excluded by our prediction scripts (**Figure 1**).

### Effect of stop codon usage on uORF repressive activity

Substitution of *HOXA11* uORF stop codon (TAG, amber) with the alternate TAA (ochre) and TGA (opal) stop codons had no significant effect on the repressive activity of the *HOXA11* uORF (**Figure S8A**). Global analysis of uORF stop codon usage revealed that UGA-terminating uORFs were the most repressive and the most common (*D*=0.135, *P*=1.45×10^−15^, *n*=1,169), compared with UAG terminating uORFs (*D*=0.0838, *P*=0.00376, *n*=483) and UAA-terminating uORFs (*D*=0.0784, *P*=0.0435, *n*=326) (**Figure S8B**).

### Effect of translation initiation site overlap on uORF repressive activity

The WT *RNASEH1* uORF overlaps with the *RNASEH1* TIS (**Figure 2A**). We generated a mutant construct (*RNASEH*1 v1) in which a stop codon and 2 additional ‘stuffer’ bases were added such that the uORF no longer overlaps with the TIS (**Figure S9A**). The non-overlapping variant was found to be repressive, although to a lesser extent than the WT construct (3.6-fold versus 5.5-fold de-repression) (**Figure S9B**). Aggregate proteomics data showed that there was no significant difference in repressive potential between uORFs which span the TIS (*n*=579) and those that are completely contained within the 5□ UTR (*n*=1,407) (*D*=0.0424, *P*=0.441. **Figure S9C**), consistent with previous observations.^1^

### Effect of amino acid composition on uORF functionality

We next sought to assess the relative contribution of each amino acid to the repressive effect of the *HOXA11* uORF. The *HOXA11* uORF peptide consists of 10 amino acids following the initial methionine (MLSSATAESPR*). An alanine scan was performed in which each single non-alanine residue was mutated to alanine in turn. In general, the effects of these mutants were minimal, with the majority exhibiting no significant effect on luciferase expression (**Figure 5A**). Substitution of the first leucine residue resulted in a 35% (*P*<0.05) reduction in reporter expression (**Figure 5A**). Mutation of any single amino acid residue had no effect on transcript levels (**Figure S10A**). Similarly, when the amino acid sequence of the uORF peptide was scrambled such that the composition remained the same, but the order was randomized, there was no significant difference in uORF-mediated translational repression relative to the WT *HOXA11* 5□ UTR (**Figure 5B**). These data suggest that individual positions in the *HOXA11* uORF contribute minimally to its repressive activity and that the order of amino acids is not a major factor affecting uORF function.

**Figure 5.**
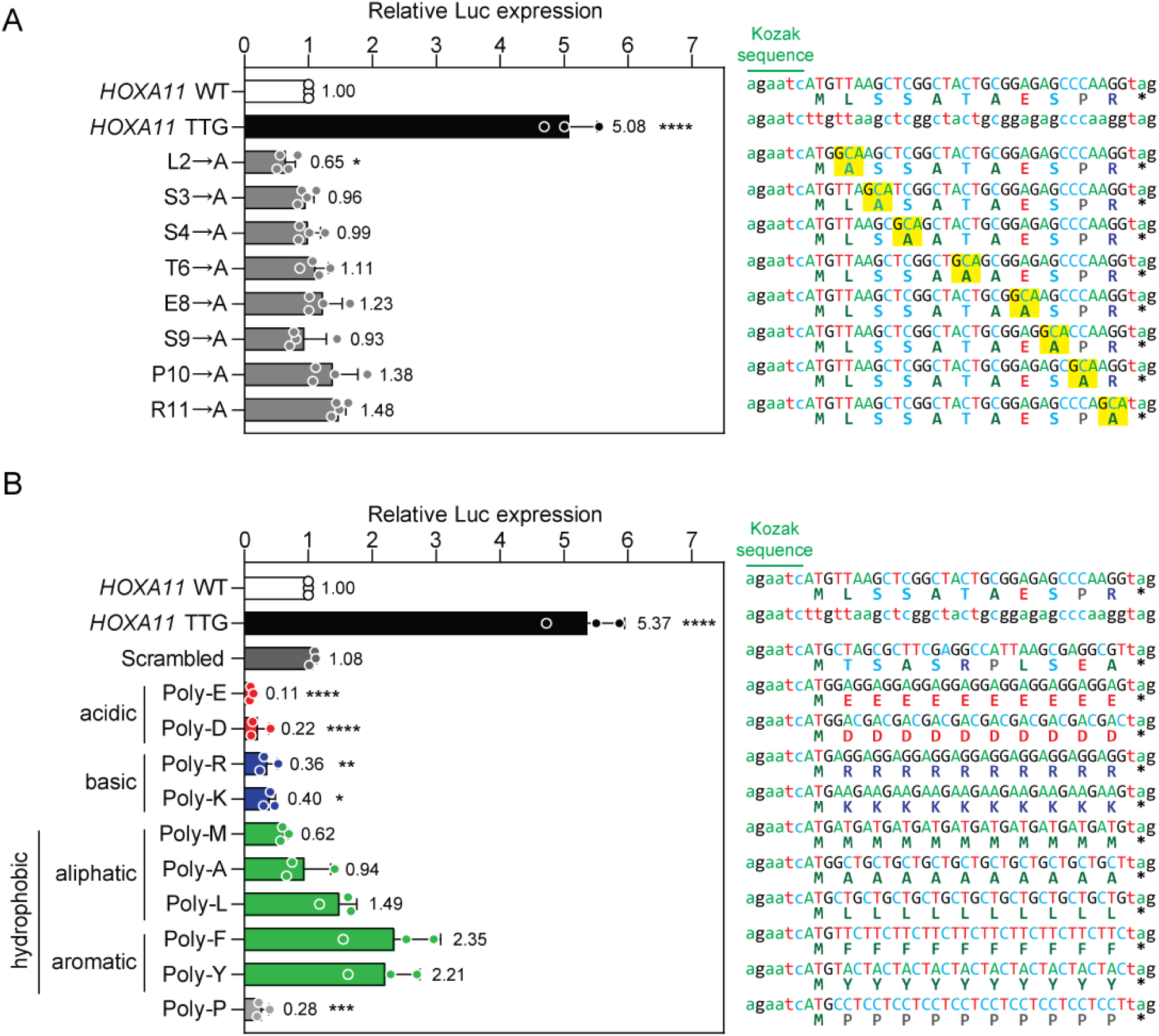
Effect of uORF amino acid composition on translation repression activity. (**A**) HEK293T cells were transfected with pDLR *HOXA11* 5□ UTR mutant vectors encoding single alanine substitutions at each non-alanine position along the *HOXA11* uORF sequence. Positions of alanine substitutions are highlighted in yellow. Dual luciferase activity was determined 24 hours post transfection. *HOXA11* WT and uORF-disrupted (TTG) constructs were included as controls. (**B**) HEK293T cells were transfected with pDLR *HOXA11* 5□ UTR mutant vectors encoding 10mer homopolymeric amino acid sequences, including acidic (E, and D), basic (R, and K), aliphatic hydrophobic (M, A, and L), aromatic hydrophobic (F, and Y), and ‘other’ (P) residues. A construct in which the *HOXA11* amino acid sequence was randomized was included as an additional control (Scrambled). Values are mean+SD (*n*=3, 4 independent experiments), and were scaled such that the mean of the WT control group was returned to a value of 1. Statistical significance was determined by one-way ANOVA with Bonferroni *post hoc* test. \**P*<0.05, \*\**P*<0.01, \*\*\**P*<0.001, \*\*\*\**P*<0.0001.

To test the effect of uORF biophysical properties on translational repression, we generated artificial uORF mutants consisting of homopolymeric amino acid sequences of ten residues long, for acidic (i.e. glutamic acid, aspartic acid), basic (i.e. arginine, lysine), aliphatic hydrophobic (i.e. methionine, alanine, leucine), aromatic hydrophobic (i.e. phenylalanine, tyrosine), and other (i.e. proline) residues. Altering uORF biophysical properties had profound effects on luciferase reporter expression. The acidic uORFs (Poly-E and Poly-D) exhibited greatly enhanced (11-22% of WT control, *P*<0.0001) repression relative to the WT control (**Figure 5B**). Similarly, the basic uORFs (Poly-R and Poly-K) also exhibited enhanced repression (36% and 40% of WT control, respectively, *P*<0.05), although to a lesser extent. In contrast, the more hydrophobic uORFs exhibited reduced translational repression, with the effect being strongest for the aromatic hydrophobic uORFs (Poly-F and Poly-Y), although these effects did not reach significance at the *P*<0.05 level (**Figure 5B**). The Poly-P construct exhibited enhanced repression (28% of WT control, *P*<0.001). The effects of the homopolymeric amino acid mutants on reporter gene activity could not be explained by changes in transcript levels as assessed by RT-qPCR (**Figure S10B).**

Further truncation mutants of selected homopolymeric uORFs were generated consisting of 1, 2, 4, 6, 8, and 10 consecutive residues of either glutamic acid (E), arginine (R), or phenylalanine (F). The enhanced repressive effects of Poly-E and Poly-R uORFs were progressively diminished as uORF length decreased (**Figure S11A,B**). Conversely, the relationship between uORF length for the poly-hydrophobic uORF, Poly-F, was less clear, although the longer Poly-F uORFs (F_6_, F_8_, and F_10_) were less repressive than the shorter Poly-F uORFs (F_1_, F_2_, and F_4_) (**Figure S11C**).

While the poly-acidic amino acid-containing uORFs exhibited enhanced repressive activity, possible confounding factors are the repetitive nature, low complexity, and high GC content of the RNA sequence encoding these artificial uORFs. To determine whether these fetaures influence uORF activity, we generated a series of additional constructs. Two codon-optimized poly-acidic uORF variants (codon optimized Poly-E incorporating multiple alternate glutamate codons, and alternating glutamate and aspartate, (ED)_5_) maintained the high level of repressive activity observed with relatively low complexity, repetitive Poly-E and Poly-D uORFs (**Figure S12A**). Mutation of the (ED)_5_ variant start codon to TTG abrogated uORF-mediated repression and demonstrated that this artificial, hyper-functional uORF represses pORF translation by ∼99% (**Figure S12B**). These results demonstrate that the repressive effects of these artificial uORFs are independent of the RNA sequence, and therefore likely a consequence of the biophysical properties of the constituent uORF amino acids.

uORF-encoding sequences containing G-quadruplexes (in two different reading frames) resulted in a small enhancement in repressive activity (5-19%), which was not significantly different from the WT *HOXA11* uORF (**Figure S12C**). Similarly, a Poly-Q uORF (encoded by (CAG)_10,_ which is expected to generate a hairpin structure),^57^ exhibited the opposite effect, whereby uORF-mediated repression was significantly reduced (**Figure S12C**). These data show that RNA secondary structure is unlikely to be responsible for the highly repressive activity of the artificial poly-acidic uORFs. Highly GC-rich uORFs encoding Poly-G (i.e. (GGA)_10_ and (GGC)_10_) exhibited enhanced repressive activity relative to the *HOXA11* WT control, but did not reach the level of repression observed for poly-acidic uORFs (**Figure S12D**), suggesting that high GC content alone cannot account for these effects.

To assess whether the availability of tRNAs for rare codons could impact uORF repressive activity we generated homopolymeric artificial uORF constructs for valine (Poly-V) and glutamine (Poly-Q), consisting of rare and common codons (**Figure S13**). No significant difference was observed when comparing rare and common codon variants (**Figure S13**).

### Effect of amino acid composition on uORF functionality interrogated using a machine learning algorithm

While artificial uORFs consisting of homopolymeric sequences exert profound effects on uORF-mediated repressive activity, such sequences do not occur naturally in uORFs. Nevertheless, these findings do point to a role for the amino acid sequence in modulating the degree of uORF-mediated repression. Given that there are 20^10^ possible permutations of the HOXA11 uORF amino acid sequence, large-scale wet lab investigations of these are impractical. Instead, we utilized the Optimus 5-Prime machine learning algorithm to interrogate putative HOXA11 uORF peptide variants *in silico*.^45^ This model has been trained on a library of 260,000 random 5□ UTR sequences, each associated with a mean ribosome load (MRL) value determined by polysome profiling that reflects its degree of translational activity. The training process has generated a network of sequence features, relationships, and rules which may not be immediately obvious or easily characterized by targeted bioinformatics analyses.^45^

To this end, we generated simulated 5□ UTR sequences in which uORF amino acid sequence could be randomized. The *HOXA11* 5□ UTR was utilized, as it is short enough (88 nucleotides) to be handled by Optimus 5-Prime, and only the 30 nucleotide/10 amino acid sequence of the uORF randomized. The 5□ UTR sequence was ‘stuffed’ with ambiguous bases (i.e. ‘N’s) at the 5□ end, as the model expects an input sequence of 100 nucleotides. The resulting simulations were presented to the Optimus 5-Prime model and predicted MRL values generated as outputs (**Figure 6A**). A series of seven simulated datasets were generated with a set of sequence constraints (*n*=2 million simulated 5□ UTRs each) with distinct properties (**Figure 6B**). The endogenous *HOXA11* uORF consists of an intermediate Kozak context due to an A nucleotide at the −3 position and a T nucleotide at the +4 position. As such, unsupervised amino acid randomization could inadvertently introduce a G at the +4 position, thereby generating a strong Kozak context. Such an effect could be confounding in terms of identifying amino acid level effects on uORF repressive activity. As such, the first nucleotide was constrained to be a G (strong Kozak context) for simulations S1, S3, and S5, or an A, C, or T (intermediate Kozak context) for simulations S2, S4, and S6 (**Figure 6B**). Similarly, we reasoned that a true random pick of simulated uORF amino acids could also introduce internal start codons (and therefore additional potential uORFs). We therefore carefully constrained the generation of the simulation sets such that simulations S1 and S2 contained exactly one ATGs per simulation, S3 and S4 contained exactly two ATGs, and S5 and S6 contained exactly three ATGs (**Figure 6B**). Considering that both nucleic acid and amino acid sequences have the potential to modify uORF activity we aimed to exclude nucleic acid level effects by generating a further simulation set (S2□) which contained identical simulated amino acid sequences to S2, but with the nucleic acid sequence codon randomized. S2□ was similarly constrained to ensure an intermediate Kozak context and to contain exactly one ATG (**Figure 6B**). In all cases, simulations were additionally constrained to prevent occurrences of additional stop codons (in-frame relative to the first ATG), as this would alter uORF length resulting in another potentially confounding factor.

**Figure 6.**
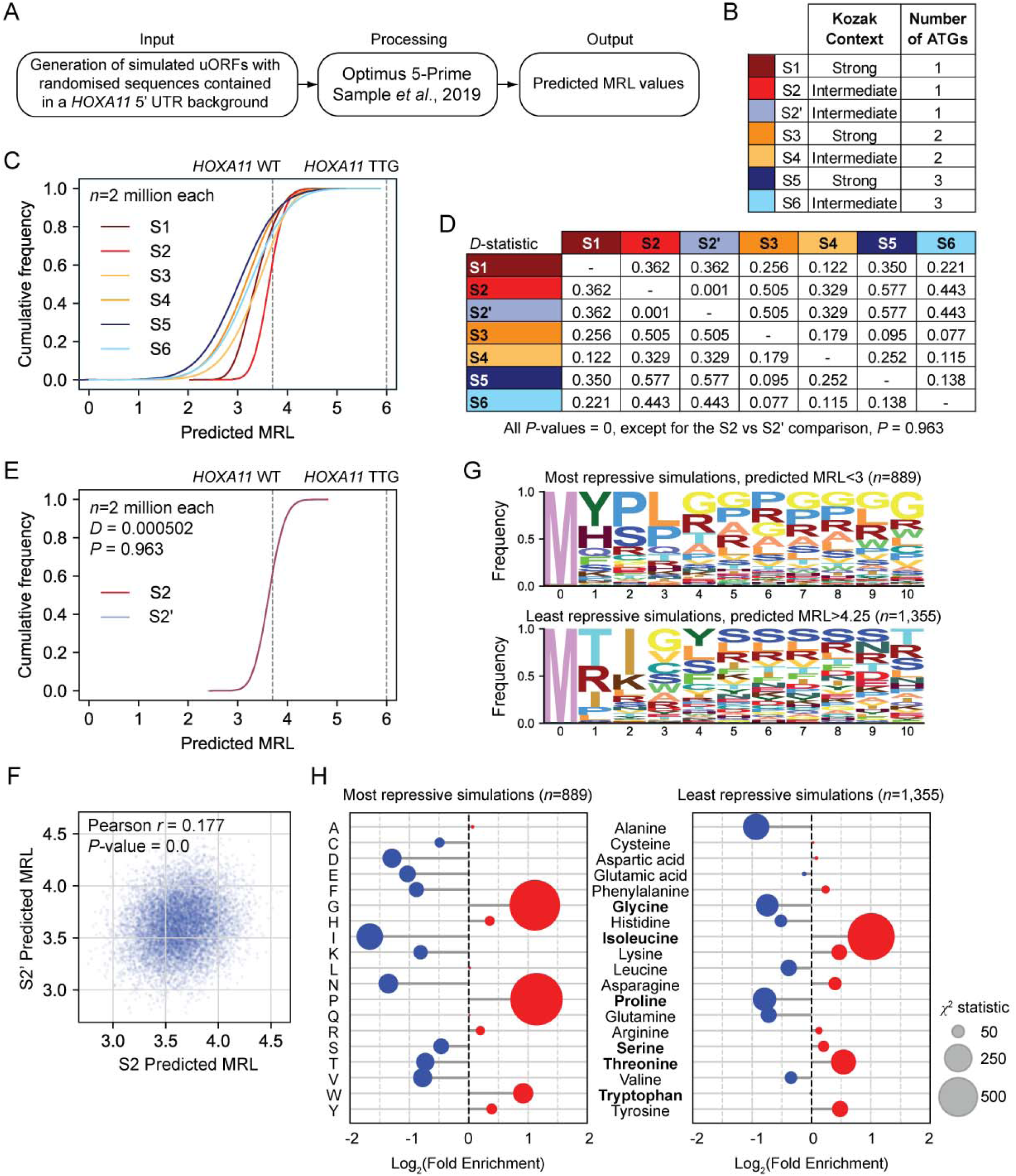
Interrogation of the effect of amino acid composition on uORF repressive activity using a machine learning approach. (**A**) Schematic of approach. Sets of simulated 5□ UTRs (2 million each) based on *HOXA11* were generated in which the amino acid sequence of the uORF was randomized. These simulations were presented to the Optimus 5-Prime algorithm, and predicted mean ribosome load (MRL) values were generated. (**B**) Table of simulation set properties (S1-S6) showing number of total ATGs and the Kozak context for the first ATG. The S2− set consists of identical amino acid sequences to S2, but with the nucleic acid sequence altered by randomized synonymous codon substitution. (**C**) CDF plots for predicted MRL values for the S1-S6 simulation sets. Predicted MRL values for *HOXA11* WT and uORF-disrupted (TTG) simulations are indicated with vertical dotted lines. (**D**) Table of pairwise Kolmogorov-Smirnov *D*-statistics. (**E**) CDF plot for S2 and S2− (synonymous codon randomized) simulation sets. Differences between the distributions were negligible and not statistically significant. (**F**) Scatter plot comparing predicted MRL values for the S2 and S2− simulation sets. (**G**) Sequence logos of enriched amino acid motifs for the most repressive simulations (mean MRL<3, *n*=889), and the least repressive simulations (mean MRL>4.25, *n*=1,355). (**H**) Lollipop plots showing enriched amino acids in the most repressive and least repressive simulated uORFs, assessed by χ^2^ test.

Analysis of the WT *HOXA11* 5□ UTR and the start codon-disrupted TTG equivalent revealed MRL values of 3.6 and 6, respectively, suggesting that the Optimus 5-Prime model has learned that the presence of a uORF reduces translation (**Figure 6C**). Analyses of CDF plots for the simulated sets similarly showed that the model has learned about other important uORF features. For example, CDFs for the strong Kozak simulations were shifted to lower MRL values relative to their intermediate Kozak context equivalents in each case, indicative of a higher probability of uORF translation initiation, and therefore repression: *D*-statistics were 0.362, 0.179, and 0.138 for strong versus intermediate Kozak contexts for the one uORF (S1 versus S2), two uORF (S3 versus S4), and three uORF (S5 versus S6) simulation sets, respectively (**Figure 6C,D**).

Similarly, increasing the number of upstream ATGs (and therefore potential uORFs) per 5□ UTR also resulted in a shift in the CDF plots towards lower predicted MRL values (*D*-statistics were 0.329 and 0.443 for one versus two ATGs (S4 versus S2) and one versus three ATGs (S6 versus S2), respectively), suggestive of an additive repressive effect of uORFs in series (**Figure S6C,D**).

We also investigated the effect of changing the identity of the initial ATG to a non-canonical start codon. To this end, variations on the S1 (strong Kozak context) and S2 (intermediate Kozak context) were generated in which the start codon was substituted for CTG, GTG, or TTG and simulated MRL values obtained (**Figure S14**). CDF plots of these simulations showed that for the strong Kozak context simulations there was a small shift in the CTG simulation set when in a strong Kozak context (S1-CTG, **Figure S14A,C**) but not in the intermediate Kozak context (S2-CTG, **Figure S14B,C**). The GTG and TTG simulation sets were otherwise very similar to the TTG sets (**Figure S14**). These data suggest that the model has also learned that translation can start at uORFs that start with a CTG trinucleotide when the Kozak context is strong, but at a rate much lower than for ATG.

These data show that the Optimus 5-Prime algorithm has learned features that influence uORF repressive activity. To investigate the impact of uORF amino acid sequence we next compared the S2 and S2− simulated datasets (**Figure 6E**). The CDF plots for these simulations were highly overlapping and not significantly different (*D*=0.001, *P*=0.963) (**Figure 6D,E**). The sigmoidal shape of the CDF plots indicates that there is substantial diversity between constituent simulations, suggesting that the model has identified features leading to their differential repressive activity. We next compared the predicted MRL for individual simulations in the S2 and S2□ simulation sets (**Figure 6F**). Surprisingly, these were only weakly positively correlated (Pearson’s *r*=0.177, *P*=0) suggesting that while amino acid composition is affecting uORF activity, there is also a substantial contribution from the nucleic acid sequence itself. Predicted MRLs for S2 and S2□ were averaged and re-ranked in order to identify those simulations that were consistently more or less repressive as a consequence of amino acid composition. Top and bottom ranking simulations were analysed using mean predicted MRL thresholds of <3 and >4.25, respectively (**Figure 6E**). Logo analysis in the most repressive simulations (*n*=889) revealed an overrepresentation of glycine and proline (**Figure 6G**). Furthermore, a prominent ‘YPL’ motif was detected at the first three amino acid positions following the methionine. Conversely, logo analysis for the least repressive simulations (*n*=1,355) revealed an overrepresentation of serine and leucine (**Figure 6G**). The prominent ‘TIGY’ motif was detected at the first four amino acid positions following the methionine. Interestingly, arginine residues were enriched in both logo analyses (**Figure 6G**). Similarly, overrepresentation of specific amino acids was analysed by χ^2^ test, which revealed that glycine, histidine, and proline were overrepresented in the most repressive simulations and underrepresented in the least repressive simulations (*P*<1.99×10^−6^, **Figure 6H**). Conversely, phenylalanine, isoleucine, asparagine, serine, and threonine were all enriched in the least repressive simulations and underrepresented in the most repressive simulations (*P*<1.76×10^−4^, **Figure 6H**).

The top ten most repressive and least repressive simulations are shown in **Figure 7A,B**. To validate machine learning predictions, we generated DLR constructs in which the native *HOXA11* uORF was substituted for artificial uORFs predicted by Optimus 5-Prime, or mutant variations thereof.

**Figure 7.**
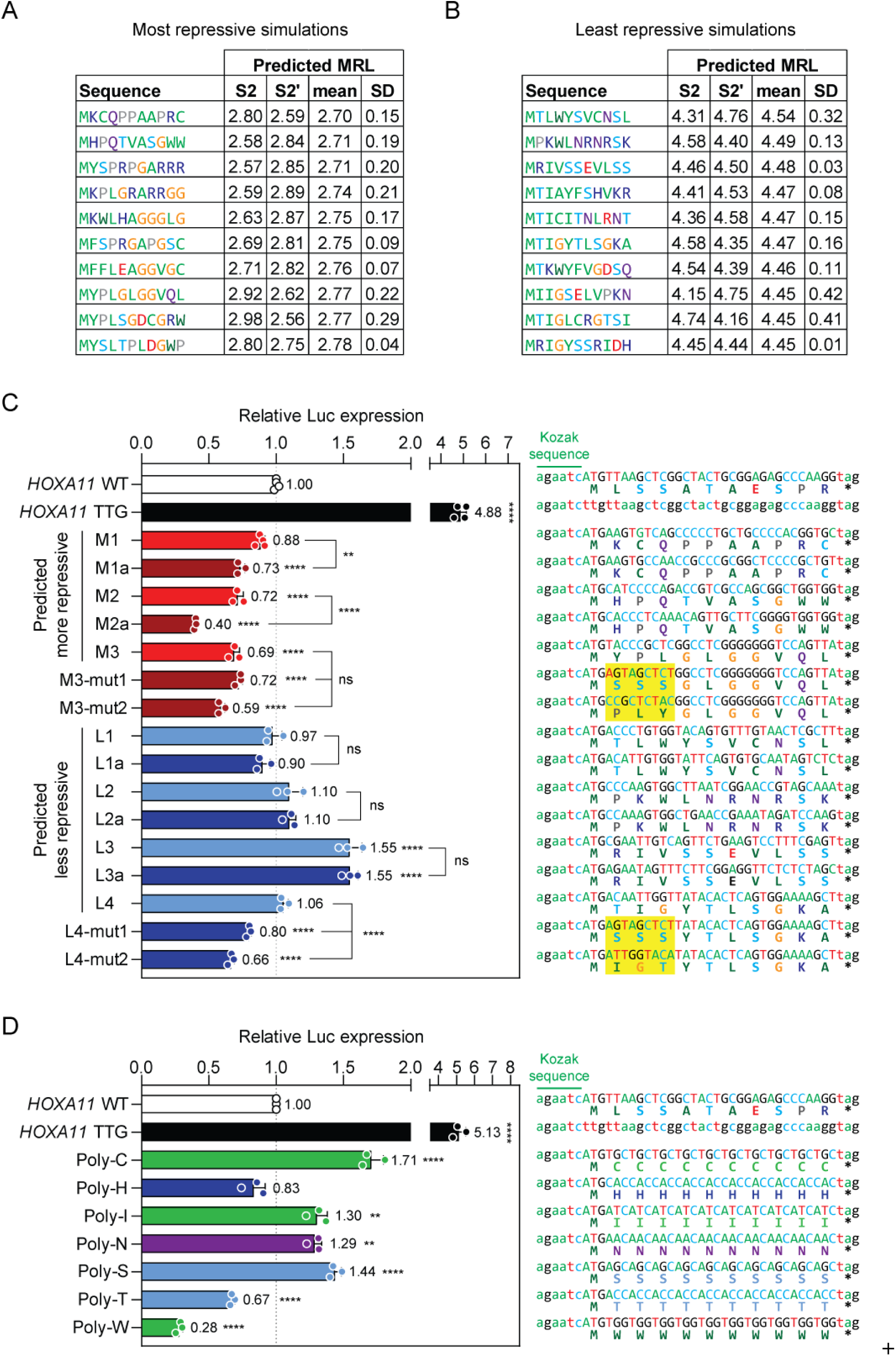
Identification of uORF activity-modulating amino acid sequences from machine learning predictions. Tables of the simulation results for (**A**) the most repressive amino acid sequences, and (**B**) the least repressive. (**C**) HEK293T cells were transfected with pDLR *HOXA11* 5□ UTR mutant vectors in which the native uORF was substituted for artificial uORFs predicted to be more, or less, repressive than the WT *HOXA11* uORF by the Optimus 5-Prime algorithm. Codon-synonymous, sequence randomized variant constructs were included as indicated. Additional mutant constructs were included in which specific motifs were disrupted (highlighted in yellow). Dual luciferase activity was determined 24 hours post transfection. *HOXA11* WT and uORF-disrupted (TTG) constructs were included as controls. (**D**) HEK293T cells were transfected with pDLR *HOXA11* 5□ UTR mutant vectors encoding 10mer homopolymeric amino acid sequences consisting of repeating residues of cysteine (C), histidine (H), isoleucine (I), asparagine (N), serine (S), threonine (T), or tryptophan (W). Values are mean+SD (*n*=3 or 4 independent experiments), and were scaled such that the mean of the WT control group was returned to a value of 1. Statistical significance was determined by one-way ANOVA with Bonferroni *post hoc* test. \*\**P*<0.01, \*\*\*\**P*<0.0001, ns=not significant.

Two uORFs M1 and M2 (each with a synonymous codon randomized construct variants; M1a and M2a) were found to be more repressive than the WT *HOXA11* uORF, consistent with machine learning predictions (**Figure 7C**). A third ‘predicted more repressive’ uORF (M3), which contained a YPL motif, was also experimentally validated as being statistically more repressive (*P*<0.0001) than the WT *HOXA11* uORF, although disruption of the YPL motif through substitution with serine residues, or scrambling the amino acid sequence, did not abrogate this effect (**Figure 7C**).

The two uORFs that were predicted to be the least repressive (L1 and L2) were not found to be statistically different from the WT *HOXA11* uORF. However, the third ‘predicted least repressive’ uORF (L3) was found to be significantly less repressive than the WT control (*P*<0.0001), consistent with the machine learning prediction. Results for the synonymous codon randomized variants (L1a, L2a, and L3a) were highly consistent to their respective partners.

A fourth ‘predicted less repressive’ uORF (L4), which contained a TIGY motif, was not statistically different from the WT *HOXA11* uORF (**Figure 7C**). However, disruption of the TIGY motif (by substitution with serine residues or scrambling the amino acid sequence) resulted in an increase in uORF repressive activity, suggesting that this motif is associated with a decrease in uORF-mediated repression (**Figure 7C**).

Considering that certain amino acids were found to be associated with changes in uORF repressive activity (**Figure 6H**), we generated artificial homopolymeric amino acid uORFs for the residues that had not been tested above (**Figure 5B, S12C, 12D, S13**). Consistent with findings from machine learning prediction, Poly-C, Poly-I, and Poly-S were statistically less repressive (*P*<0.01), whereas Poly-W was found to be statistically more repressive (*P*<0.0001), with a 72% reduction in expression compared to the WT *HOXA11* uORF (**Figure 7D**). Poly-H was found to be more repressive (reduced by 17% relative to the WT control), although this did not reach statistical significance at the *P*<0.05 level. By contrast, results for Poly-N and Poly-T did not match the machine learning predictions, although both were statistically different from the control (*P*<0.01) but in the opposite direction to that expected (**Figure 7D**).

In summary, using a combination of Optimus 5-Prime machine learning algorithm and experimental validation we were able to identify specific artificial uORF sequences with either greater or lesser repressive activity relative to the WT *HOXA11* uORF, identify a motif (i.e TIGY) associated with reduced uORF repressive activity, and identify amino acids that were associated with either greater (i.e. tryptophan and threonine) or lesser (i.e. cysteine, isoleucine, asparagine, and serine) pORF repression.

### Effect of uORF start codon position on functionality

To investigate the effect of the position of a uORF within the 5□ UTR on activity, we selected a gene with no predicted uORFs, *BRCA2*, and generated mutant constructs in which the *HOXA11* uORF sequence (and WT Kozak context) was inserted at the start, middle, or end of the *BRCA2* 5□ UTR (i.e. proximal, medial, or distal relative to the pORF TIS). Additional constructs were tested in parallel in which the uATG was mutated to TTG to disrupt uORF activity, or the *HOXA11* uORF was replaced with the hyper-functional (ED)_5_ artificial uORF sequence (**Figure S12A**). The introduction of the *HOXA11* uORF into the *BRCA2* 5□ UTR resulted in downstream pORF repression, which was strongest in the pORF proximal position (*P*<0.0001), intermediate in the medial position, and almost completely relieved in the distal position (**Figure 8A**). No repressive activity was observed with any of the corresponding TTG mutant controls. A similar positional effect was observed with the artificial (ED)_5_ uORF, with the highest repressive activity observed in the pORF proximal potion. However, the (ED)_5_ uORF was still functional (>90% reduction relative to the *BRCA2* WT control) in the distal position, suggesting that optimized uORF amino acid composition can override the effect of uORF position (**Figure 8A**). Luciferase data could not be explained by changes in mRNA levels (**Figure S15**) consistent with translation level effects.

**Figure 8.**
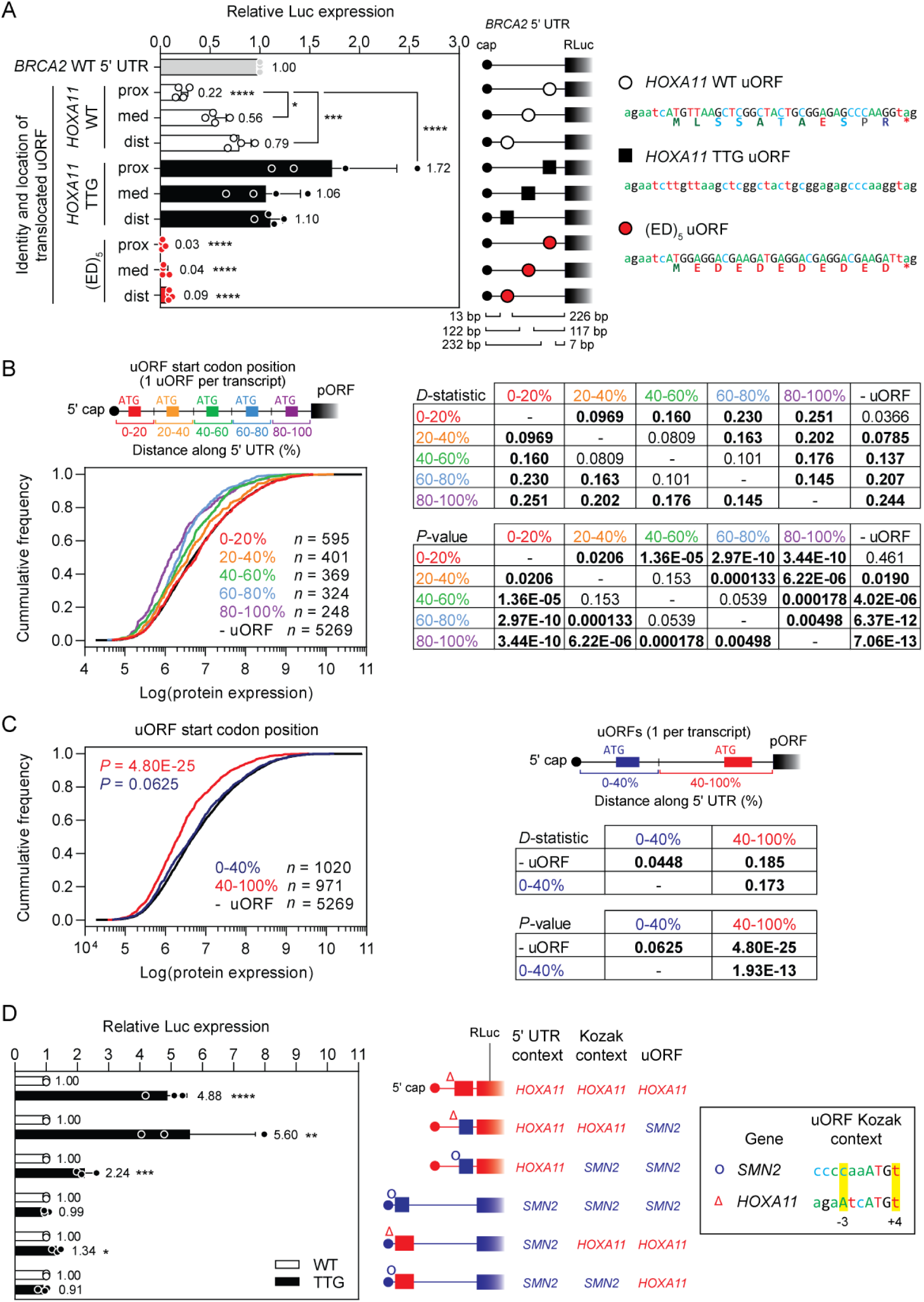
Effect of translation initiation codon position on uORF activity. A dual luciferase reporter was generated whereby the (uORF naïve) *BRCA2* 5□ UTR was cloned up-stream of the *Renilla* luciferase gene. Mutant variants were generated in which the *HOXA11* uORF (and Kozak context) were cloned into the *BRCA2* 5□ UTR at three different positions (proximal, distal, and medial relative to the pORF). Inactive versions of these constructs were generated in parallel in which the *HOXA11* uATG was mutated to TTG. Additional variants were generated in which the *HOXA11* uORF was substituted with a hyper-functional artificial (ED)_5_ uORF. (**A**) These constructs were transfected in HEK293T cells and luciferase activity assayed after 24 hours. Values are mean+SD (*n*=3 independent experiments), and were scaled such that the mean of the WT control group was returned to a value of 1. Statistical significance was determined by one-way ANOVA with Bonferroni *post hoc* test. Aggregated proteomics data from 29 healthy human tissues were binned according to the position within 5□ UTR where a predicted uORF starts and data presented as cumulative distribution function plots. Transcripts were divided into (**B**) equal 20% bins, or (**C**) a 5□ 40% bin and a 3□ 60% bin. Statistical differences between distributions were assessed by Kolmogorov-Smirnov test and *D*-statistics and *P*-values tabulated. The numbers of proteins in each bin are indicated. (**D**) Constructs were generated in which the inactive *SMN2* uORF was inserted into the *HOXA11* 5□ UTR. Conversely, the *HOXA11* uORF was inserted into the *HOXA11* 5□ UTR. In each case, both the *SMN2* and *HOXA11* uORF Kozak contexts were tested. The sequences of the two Kozak contexts are indicated. The native *HOXA11* and *SMN2* 5□ UTRs were included as controls, together with uORF-inactivated (TTG) version of each construct. Values are mean+SD (*n*=3 independent experiments). Statistical significance was determined by a Student’s *t*-test relative to the corresponding WT control. \**P*<0.05, \*\**P*<0.01, \*\*\**P*<0.001, \*\*\*\**P*<0.0001.

To assess the effects of uORF position on repressive activity for all genes, single uORF-containing transcripts were assigned to separate bins depending on the location of the uATG and CDF analysis performed using aggregated human proteomics data, as above. For this purpose, transcripts were divided into 5 bins, representing 20% increments along the length of the 5□ UTR starting at the cap. Aggregated global protein expression in the first bin, 0-20% (*n*=595) was not statistically different from genes that contained no uORFs (**Figure 8B**). However, all other bins exhibited statistically reduced protein expression CDFs relative to the no uORF bin, with the *D*-statistic becoming progressively larger (and more significant) with closer proximity to the pORF (**Figure 8B**). This effect was even more pronounced when the data were considered as only two approximately equally sized bins (0-40%, *n*=1,020 and 41-100%, *n*=971). The distribution of protein abundance values for the 0-40% bin was highly similar, although statistically different, to that of transcripts containing no uORFs (*P*=0.0625, *D*=0.0448). Conversely, the distribution for the 41-100% bin was greatly reduced (*P*=4.80×10^−25^, *D*=0.185) (**Figure 8C**).

Given the strong position-dependence of uORF activity observed with the *HOXA11* uORF, we next sought to address the question of whether a non-functional uORF could be re-activated by altering its position within the 5□ UTR. Accordingly, uORF-swapping constructs were generated in which the non-repressive *SMN2* uORF (**Figure S4**) was inserted into the *HOXA11* 5□ UTR (replacing the native uORF), and conversely, the functional *HOXA11* uORF was inserted into the *SMN2* 5□ UTR to replace the native uORF in its cap-proximal position. Both *SMN2* and *HOXA11* Kozak contexts were tested (weak and intermediate, respectively), and uORF-ablated ‘TTG’ constructs used as controls for each configuration (**Figure 8D**). The ‘non-functional’ *SMN2* uORF was observed to be functional at repressing downstream reporter expression when placed in a favorable pORF-proximal position within the *HOXA11* 5□ UTR (*P*<0.01). This effect was most pronounced when the *HOXA11* Kozak context was utilized, with the repressive effect close to that of the native *HOXA11* uORF (**Figure 8D**). Conversely, the ‘functional’ *HOXA11* uORF was found to exhibit only minimal repressive activity when placed in the unfavorable cap-proximal *SMN2* context (**Figure 8D**). Taken together, these data show that the location of the uORF start position within the 5□ UTR is a major determinant of repressive activity, with close proximity to the pORF associated with greater levels of repression.

### Effect of uORF stop codon position on functionality

The relative lack of translational repressive activity of cap-proximal uORFs might be the result of a failure of the ribosome to initiate. Metagene analysis of aggregated Ribo-Seq data across the human protein-coding transcriptome showed that mean read density was lowest in the first 20% of the 5□ UTR for both initiating ribosome peaks (**Figure 9A**) and ribosome footprints (**Figure 9B**). These data suggest that ribosome initiation at cap-proximal uORFs is possible but less likely than at other positions, consistent with previous reports.^58,59^

**Figure 9.**
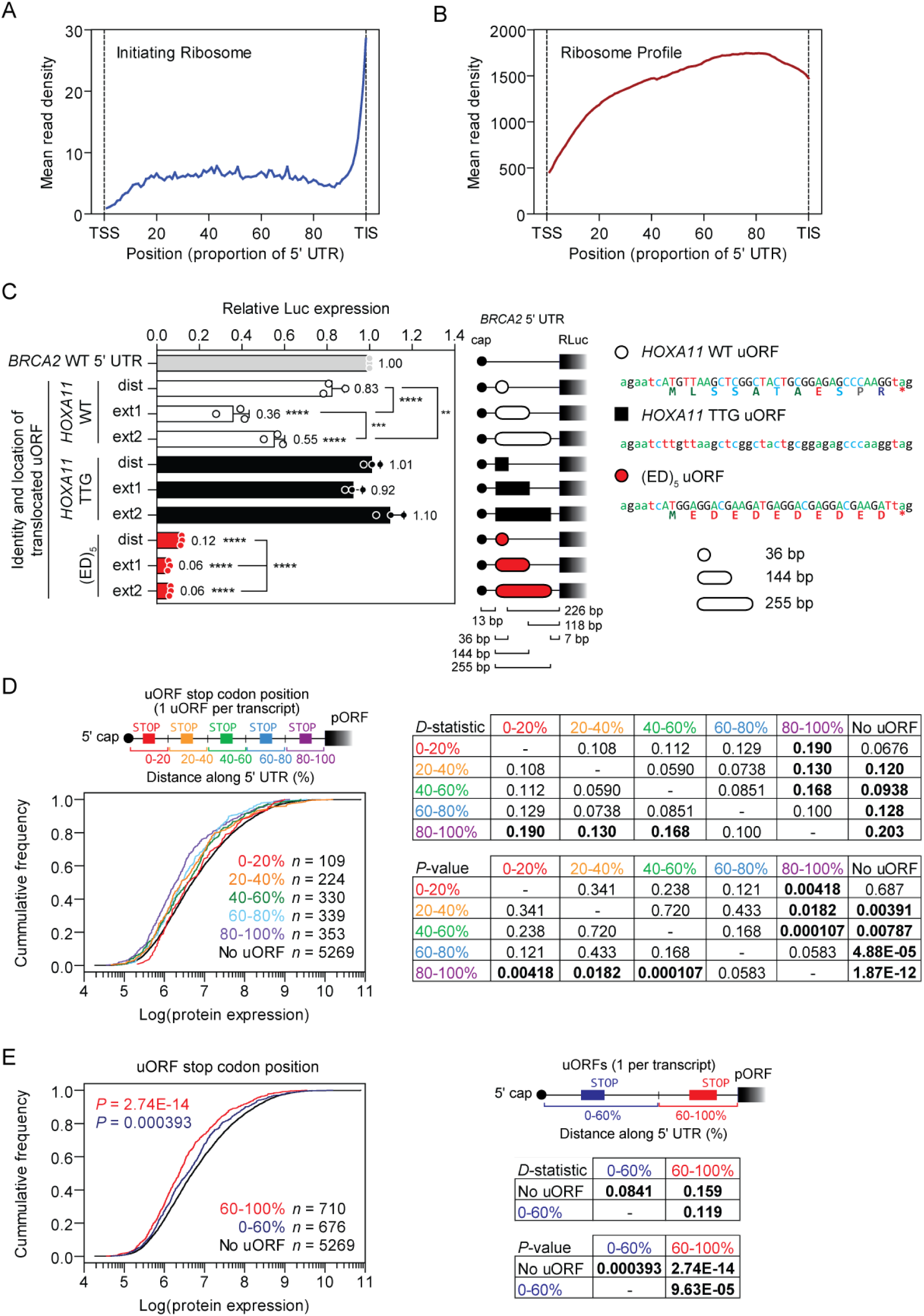
Effect of translation termination codon position on uORF activity. Metagene analysis of aggregated Ribo-Seq data for (**A**) initiating ribosome peaks, and (**B**) ribosome footprints. Plasmid DNA constructs were generated in which the *HOXA11* uORF (and Kozak consensus sequence) were cloned into the *BRCA2* 5□ UTR at the pORF distal position relative to the pORF and the 36 bp uORF extended to either 144 or 255 bp (ext1 and ext2, respectively) by shifting the position of the uORF stop codon. Inactive versions of these constructs were generated in parallel in which the *HOXA11* uATG was mutated to TTG. Additional variants were generated in which the *HOXA11* uORF was substituted with a hyper-functional artificial (ED)_5_ uORF. (**C**) These constructs were transfected in HEK293T cells and luciferase activity assayed after 24 hours. Values are mean+SD (*n*=3 independent experiments), and were scaled such that the mean of the WT control group was returned to a value of 1. Statistical significance was determined by one-way ANOVA with Bonferroni *post hoc* test, \*\**P*<0.01, \*\*\**P*<0.001, \*\*\*\**P*<0.0001. Aggregated proteomics data from 29 healthy human tissues were binned according to the position within 5□ UTR where a predicted uORF stops and data presented as cumulative distribution function plots. Transcripts were divided into (**D**) equal 20% bins, or (**E**) a 5□ 60% bin and a 3□ 40% bin. Statistical differences between distributions were assessed by Kolmogorov-Smirnov test and *D*-statistics and *P*-values tabulated. The numbers of proteins in each bin are indicated.

We next reasoned that the position of the uORF stop codon might similarly affect repressive activity. uORFs that initiate close to the pORF must necessarily also have stop codons that are close to, or overlapping with, the pORF. We therefore designed an experiment to deconvolute the effects of uORF start and stop position. The *BRCA2* 5□ UTR was again utilized as a uORF-free background in which the *HOXA11* uORF or artificial (ED)_5_ uORF was inserted in cap-proximal positions. While these start codons are close to the cap (i.e. pORF distal), our previous data illustrating the repressive activity of the (ED)_5_ uORF in this position strongly suggest that translation initiation can indeed occur at this position (**Figure 8A, S12B**). Plasmid constructs were generated in which these uORFs were artificially extended by shifting their stop codons to positions at the center of the 5□ UTR (ext1) and proximal to the pORF (ext2). Both stop codon extension constructs resulted in a reduction in relative luciferase reporter expression, which was more pronounced for the ext1 construct (36% of WT, *P*<0.0001) (**Figure 9C**). A similar effect was observed for the artificial (ED)_5_ uORFs, whereby extension of these uORFs resulted in an increase in repressive activity. No repressive activity was observed with any of the corresponding TTG mutant controls (**Figure 9C**). Luciferase data could not be explained by changes in mRNA levels (**Figure S16**) consistent with translation level effects.

To assess the effects of uORF stop position on repressive activity for all genes, single uORF-containing transcripts were assigned to separate bins depending on the location of the uORF termination codon. For this purpose, transcripts were divided into 5 bins, representing 20% increments along the length of the 5□ UTR starting at the cap. Aggregated global protein expression in the first bin, 0-20% (*n*=109) was not statistically different from genes that contained no uORFs (**Figure 9D**). However, all other bins exhibited statistically reduced protein expression CDFs relative to the no uORF bin with the effect becoming more significant with closer proximity to the pORF (**Figure 9D**).

Transcripts were reclassified into two bins of approximately equal size (0-60% distance, *n*=676, and 60-100% distance, *n*=710) and reanalyzed in order to maximize statistical power. Termination codons located in the later region (60-100%) of the 5□ UTR exhibited the greatest shift in CDF curve relative to the No uORF bin (*D*=0.159, *P*=2.74×10^−14^) (**Figure 9E**). By contrast, for the earlier region (0-60%), a CDF curve shift relative to the no uORF bin was observed, although with lower magnitude and statistical significance (*D*=0.0841, *P*=0.000393) (**Figure 9E**).

We next sought to determine whether the activity of an endogenous uORF could be modulated by altering the position of its stop codon. We selected *CUL2* (NM_003591.4) as a candidate 5□ UTR as it contains a single, short cap-proximal uORF (starting at position 15) for which there was evidence of translation from aggregated Ribo-Seq data (**Figure 10A**). While an initiating ribosome peak was observed at the predicted *CUL2* uORF, its peak height was much less than that observed at the *CUL2* pORF (uORF:pORF initiating ribosome peak height ratio: 0.22) (**Figure 10A**). These data suggest that while the *CUL2* uORF is translated, it is unlikely to be suppressing translation of the downstream *CUL2* pORF.

**Figure 10.**
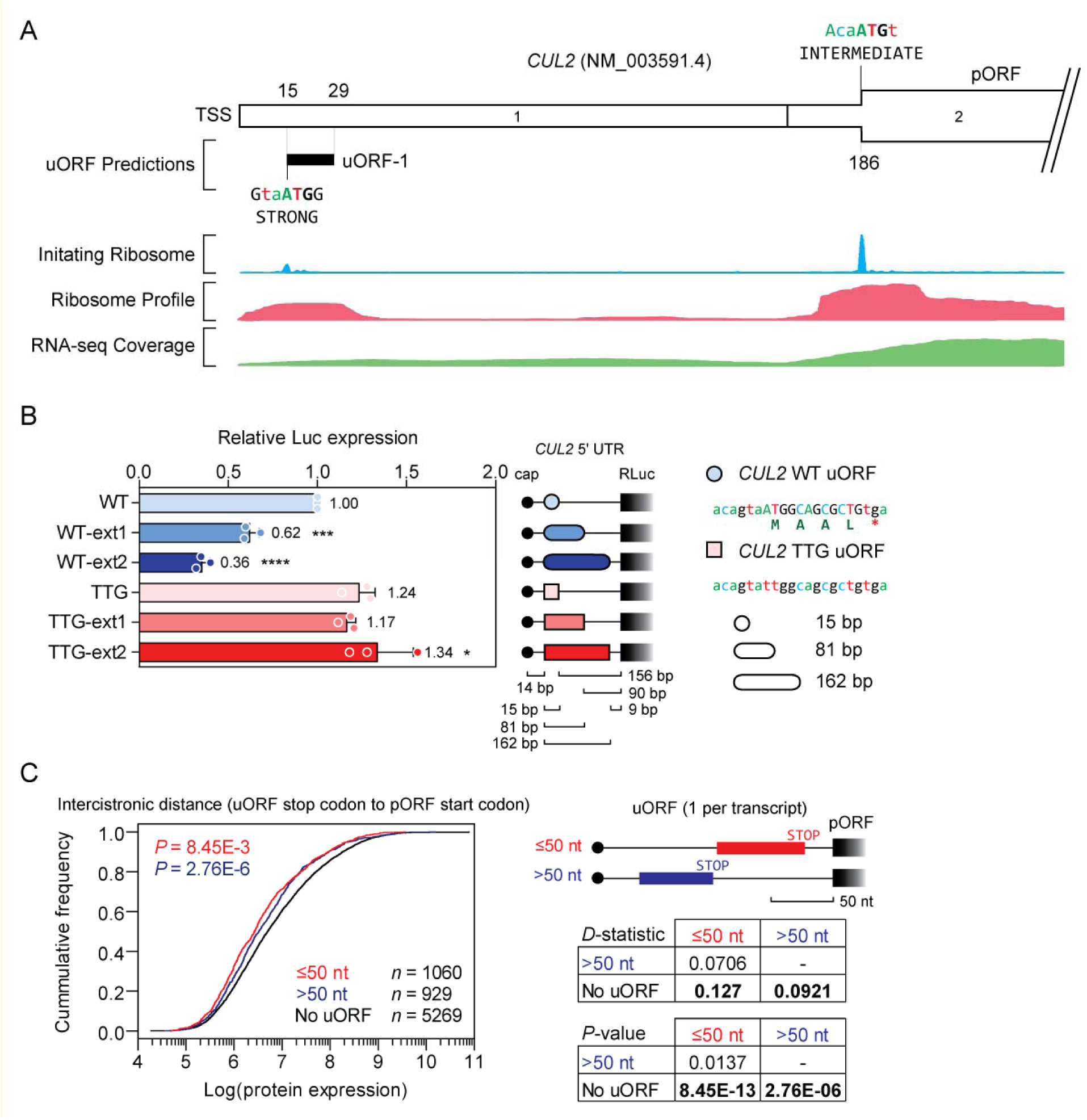
Effect of intercistronic distance on uORF activity. (**A**) Schematic of *CUL2* transcript variant NM_003591.4, containing 1 predicted uORF located distal to the pORF. Aggregated Ribo-Seq and RNA-Seq data are overlaid which suggest that the *CUL2* uORF is translated but exerts minimal repressive effects on the pORF. (**B**) HEK293T cells were transfected with *CUL2* 5□ UTR dual luciferase reporter constructs as indicated and luciferase activity assayed after 24 hours. Two variant constructs were generated in which the 15 bp *CUL2* uORF was extended (WT-ext1 and WT-ext2). Corresponding TTG variants (in which the uORF start codon was disrupted) were generated in each case. Values are mean+SD (*n*=3 independent experiments), and were scaled such that the mean of the WT control group was returned to a value of 1. Statistical significance was determined by one-way ANOVA with Bonferroni *post hoc* test. (**C**) Aggregated proteomics data from 29 healthy human tissues were binned according to the position within 5□ UTR where a predicted uORF ends and data presented as cumulative distribution function plots. Transcripts were binned according to whether the intercistronic distance between the uORF and pORF was greater than, or less than, 50 nucleotides. Statistical differences between distributions were assessed by Kolmogorov-Smirnov test and *D*-statistics and *P*-values tabulated. The numbers of proteins in each bin are indicated. \**P*<0.05, \*\*\**P*<0.001, \*\*\*\**P*<0.0001.

Dual luciferase reporter constructs were generated in which the *CUL2* 5□ UTR was cloned upstream of Renilla luciferase. The intercistronic distance between the WT termination codon and the pORF start codon is 156 bp. Two mutant constructs (WT-ext1 and WT-ext2) were generated in which the uORF was progressively extended such that the intercistronic distances were 90 and 9 bp, respectively. Control constructs in which the uORF was disrupted by mutating the start codon from ATG to TTG were tested in parallel. Consistent with the Ribo-Seq-driven predictions (**Figure 10A**), the WT *CUL2* uORF was found to exhibit negligible repressive activity. However, extension of the uORF to reduce the intercistronic distance progressively repressed the downstream reporter relative to the WT construct, or to any of the TTG controls (**Figure 10B**). These data show that a shift in the termination codon position (and therefore also intercistronic distance) can convert a largely inactive uORF into an active uORF.

To investigate the impact of inter-cistronic distance on a global scale, single uORF-containing transcripts were assigned to separate bins depending on their distance (in nucleotides) relative to the pORF. Two bins, ≤50 nt or >50 nt, of approximately equal sample size (*n*=1,060 and *n*=929, respectively) were selected for reasons of statistical power, and because this distance corresponds to a value in the middle of the proposed range for the refractory period required for effective reinitiation.^36^ uORFs that terminated within 50 nt of the pORF exhibited a greater CDF curve shift (*D*=0.127, *P*=8.45×10^−13^) than uORFs terminating >50 nt from the pORF (*D*=0.0921, *P*=2.76×10^−6^) (**Figure 10C**). Taken together, these data show that the location of the uORF termination codon within the 5□ UTR is a major determinant of repressive activity, with close proximity to the pORF being associated with greater levels of repression.

### Interaction between uORFs in multi-uORF contexts

We next sought to investigate transcripts containing multiple uORFs. Analysis of aggregated proteomics data revealed that transcripts with >1 uORF tended to be more repressed than those with 1 uORF, suggesting that genes with multiple uORFs are subject to a greater degree of uORF-mediated repression (**Figure 11A**). Two multi-uORF-containing transcripts were selected for detailed analysis: *BRD3* (NM_007371.4, two predicted uORFs) and *ATF4* (NM_182810.3, three predicted uORFs).

**Figure 11.**
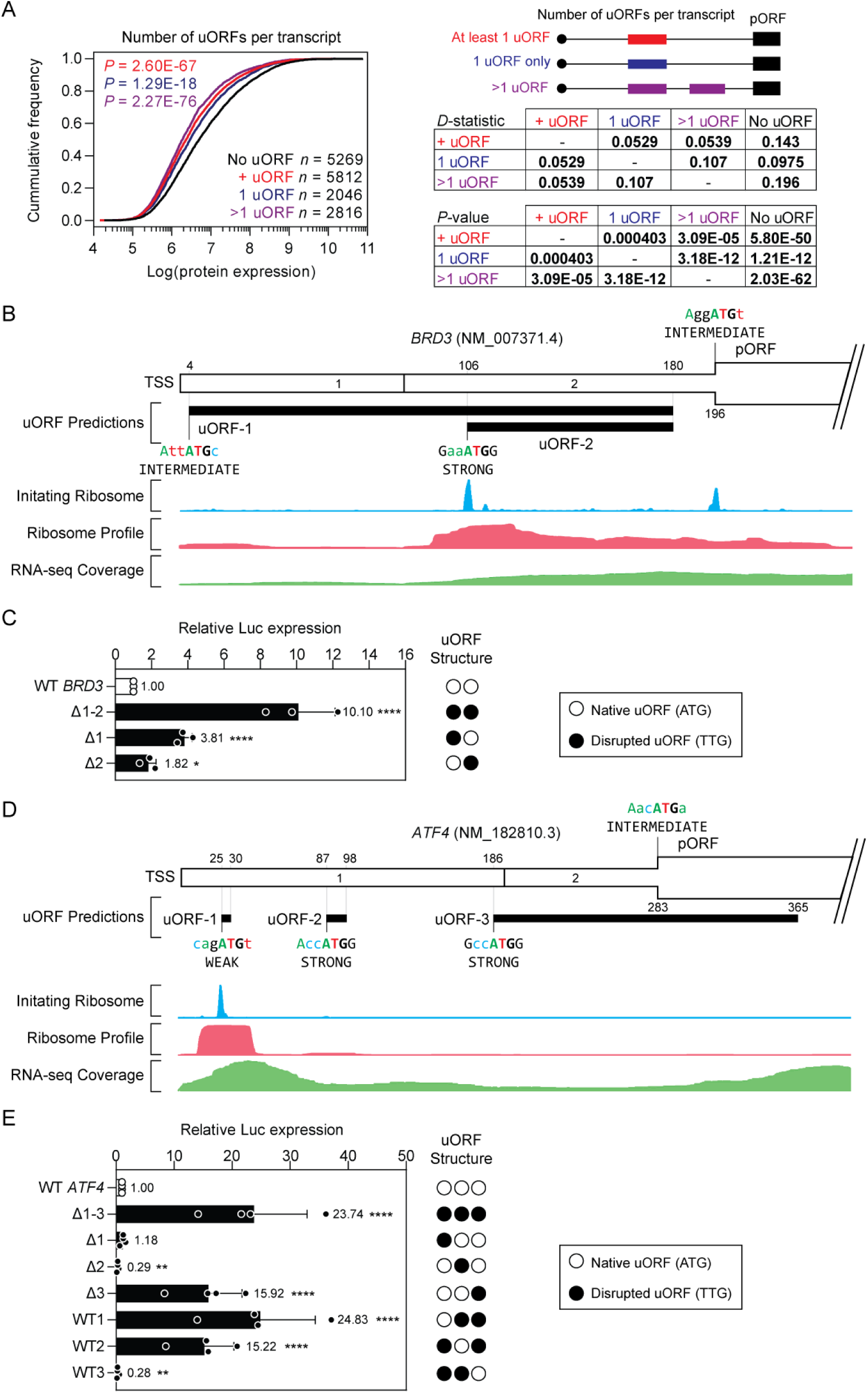
Effect of multiple uORFs on translational repression. (**A**) Aggregated proteomics data from 29 healthy human tissues were binned according to the number of predicted uORFs and data presented as cumulative distribution function plots. Transcripts were binned according to whether they contained at least 1 uORF, exactly 1 uORF, or more than 1 uORF. Statistical differences between distributions were assessed by Kolmogorov-Smirnov test and *D*-statistics and *P*-values tabulated. The numbers of proteins in each bin are indicated. (**B**) Schematic of *BRD3* transcript variant NM_007371.4, containing 2 predicted uORFs. Aggregated Ribo-Seq and RNA-Seq data are overlaid. (**C**) HEK293T cells were transfected with *BRD3* 5□ UTR dual luciferase reporter constructs and luciferase activity assayed after 24 hours. Mutant constructs in which one or both of the predicted uORFs were disrupted were analysed in parallel. (**D**) Schematic of *ATF4* transcript variant NM_182810.3, containing 3 predicted uORFs. Aggregated Ribo-Seq and RNA-Seq data are overlaid. (**E**) HEK293T cells were transfected with *ATF4* 5□ UTR dual luciferase reporter constructs and luciferase activity assayed after 24 hours. Mutant constructs in which the predicted uORFs were disrupted in variations as indicated were analysed in parallel. Values are mean+SD (*n*=3 or 4 independent experiments), and were scaled such that the mean of the WT control group was returned to a value of 1. Statistical significance was determined by one-way ANOVA and Bonferroni *post hoc* test, \**P*<0.05, \*\**P*<0.01, \*\*\*\**P*<0.0001.

Analysis of *BRD3* revealed three predicted uORFs, with prominent Ribo-Seq initiation peaks and footprints at the second uORF, although sequencing coverage was overall lower over the first 5□ UTR, which contained the first predicted uORF (**Figure 11B**). The *BRD3* 5□ UTR was cloned into the dual luciferase reporter system and mutant constructs were generated in which the uORFs in these 5□ UTRs were disrupted by ATG-to-TTG substitutions, in various combinations as indicated. Mutation of both uORFs in the *BRD3* 5□ UTR resulted in a ∼10-fold increase in reporter gene expression (*P*<0.0001, **Figure 11C**). Disruption of either uORF alone resulted in statistically significant increases in reporter expression, although these did not reach the level observed for disruption of all uORFs (**Figure 11C**). These data demonstrate that both BRD3 uORFs are functional and work synergistically to repress downstream pORF expression. Both uORF-1 and uORF-2 are in-frame with one another, sharing a stop codon that is very close to the pORF, a configuration consistent with high uORF-mediated repression (**Figures 9-10**).

*ATF4* has been extensively studied in the context of uORF-mediated regulation, ^37–40,60,61^ and contains three uORFs (**Figure 11D**). The first (minimal) uORF was associated with a prominent ribosome footprint and translation initiation peak, whereas ribosome coverage on the other uORFs was negligible (**Figure 11D**). Removal of all three uORFs resulted in a ∼24-fold upregulation of downstream reporter expression (*P*<0.0001, **Figure 11E**). This effect could mostly be attributed to uORF-3 (which overlaps with the pORF), whereby a ∼16-fold upregulation was observed when this single uORF was disrupted in isolation (*P*<0.0001, **Figure 11E**). Interestingly, when uORF-2 was disrupted, pORF repression was enhanced (∼80% reduction relative to the WT *ATF4* 5□ UTR, *P*<0.01) **(Figure 11E**). These data are consistent with a model in which uORFs interact, such that the proximity between uORF-2 and uORF-3 results in the former constraining the repressive activity of the latter.^37^ uORF-1 is a cap-proximal uORF that terminates far (252 nt) from the pORF, which may explain its relative lack of repressive activity. Taken together, these data show that a variety of complex phenomena occur in the case of 5□ UTRs with multi-uORF architectures.

## Discussion

Here we have predicted occurrences of uORFs in the human and mouse transcriptomes (**Figure 1**) and investigated the relationship between uORF features and their potential to influence the expression of their corresponding downstream pORFs. A panel of candidate 5□ UTRs/uORFs was selected based on potential disease relevance in cancer biology, of which the majority were experimentally validated as being pORF-repressive (**Figure 2**). Results for positive control uORFs: *KDR*, *MAP2K2*, and *RNASEH1*, were similar to those reported previously.^1,28,62^ Conversely, we did not observe any significant translational repression for the *SRY* uORF (which was reported to be highly repressive by others)^1^ (**Figure 2B**). Importantly, certain uORFs displayed negligible repressive activity (e.g. *SMN2* and *UTRN*; **Figure S4**). These findings underscore the importance of experimental validation in uORF research and point to possible context-dependent factors which may be influencing uORF function.

To investigate how uORF features influence their activity, we employed a combined experimental and bioinformatic approach. We primarily focused on the *HOXA11* tumor suppressor gene as a model system in which to experimentally manipulate uORF properties by generating a wide variety of mutant constructs. This relatively simple gene was selected as it is readily amenable to experimental manipulation and analysis using Optimus 5-Prime (given the size of the *HOXA11* 5□ UTR, its single uORF, and uncomplicated transcript landscape), and given its importance in development and cancer.^63–66^ The *HOXA11* uORF exhibits archetypal properties, including ribosome occupancy (**Figure 3A**), in-*cis* pORF translational repression (**Figure 3C,D**), and was repressive when inserted into other 5□ UTR contexts (**Figures 8A,D,9C**). In parallel, we utilized matched transcriptomics/proteomics datasets from 29 healthy human tissues to bin transcripts according to their uORF properties and then plot the CDFs for protein expression.^44^ The richness of the datasets used for these analyses have allowed for higher confidence inferences and more property comparisons relative to similar approaches described previously.^1^ In this manner, the extent to which inferences from single gene studies could be applied across the transcriptome was assessed for multiple uORF features including uORF Kozak context strength, length, start/stop codon position, relative reading frame, pORF overlap, and the presence of multiple uORFs. Given the diversity of possible uORF peptide sequences, we were unable to perform a meaningful CDF-type analysis to investigate amino acid composition on a wider scale. Instead, we utilized a published machine learning algorithm^45^ to predict the translation efficiency of two million randomly-generated uORFs that replaced the WT uORF embedded within the *HOXA11* 5□ UTR.

Based on these studies, an overall model of uORF-mediated translational repression emerges (**Figure 12**). The 40S small ribosomal subunit binds at the m^7^G 5□ cap and scans through the 5□ UTR until a suitable initiation codon is reached, at which point the 80S ribosome assembles and translation initiation occurs.^67–70^ For transcripts lacking uORFs (**Figure 12A**), this is at the pORF start codon. Alternatively, in situations where a uORF is present in the 5□ UTR, translation initiation can occur at the upstream start codon. Upon translation termination at the uORF stop codon, the 80S ribosome may fully dissociate from the mRNA (i.e. ribosome recycling),^71,72^ meaning that the pORF is not translated with respect to the ribosome in question (**Figure 12B**). In some cases, the 40S subunit remains associated with the transcript after uORF translation termination and continues scanning through the mRNA transcript.^8–10^ If a second uORF is present, the uORF translation process repeats, potentially leading to a second ribosome recycling event (**Figure 12C**). As the frequency of ribosome recycling events in a 5□ UTR increases, the degree of pORF repression would be expected to increase accordingly, as fewer translation-competent ribosomes reach the pORF start codon. This notion is supported by the observation that transcripts containing multiple uORFs tend to be more repressed (**Figure 11A**), consistent with previous observations.^41,73^ Such an eventuality is clearly exemplified by *BRD3*, in which the repressive effect of its two uORFs is additive (**Figure 11B,C**). Importantly, transcripts with multi-uORF configurations constitute the majority of all predicted uORF-containing transcripts (human: 65.6%, mouse: 59.2%; **Figure 1B**).

**Figure 12.**
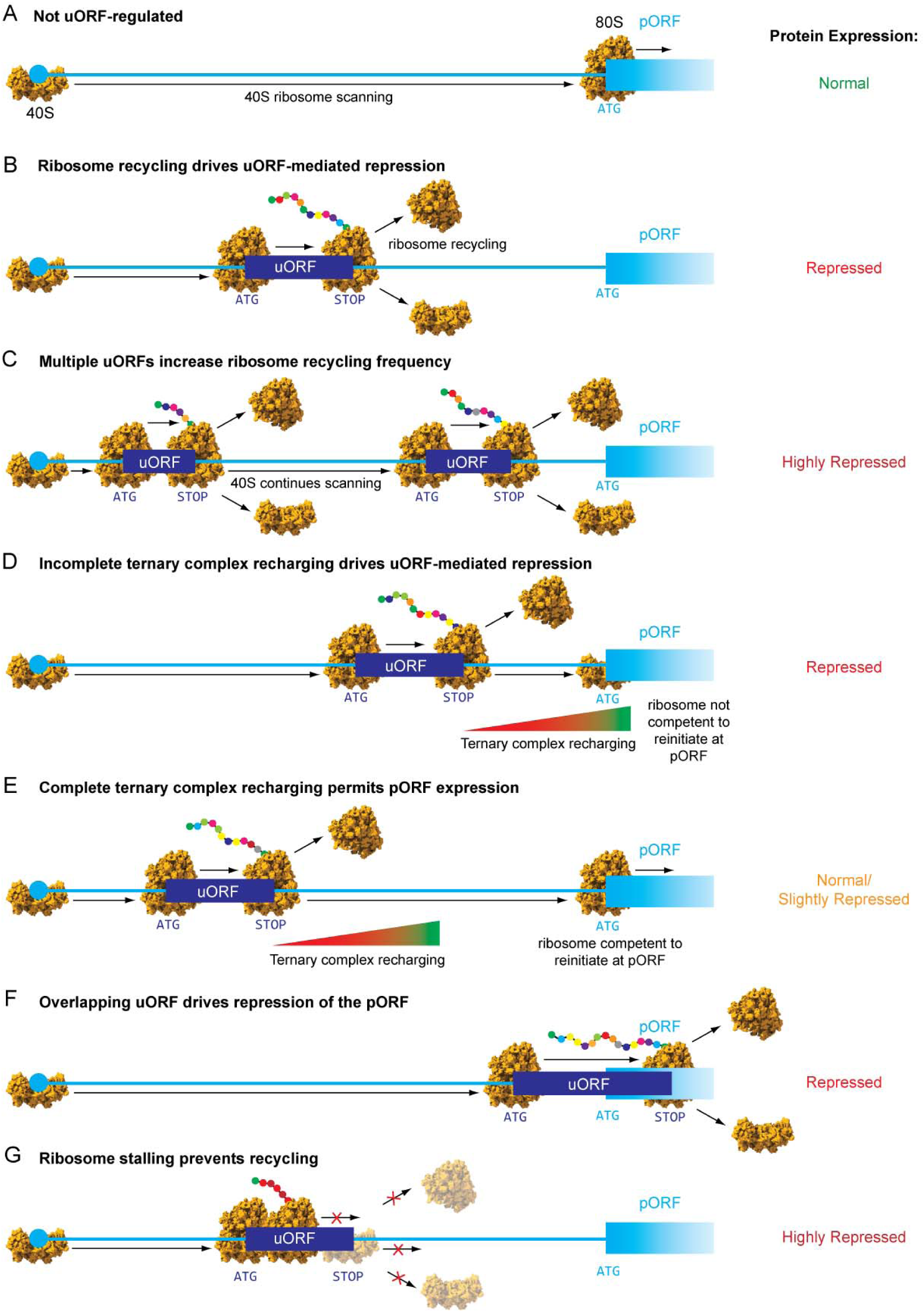
Models of uORF-mediated translational repression. (**A**) Cartoon schematics showing a transcript that is not uORF regulated. The 40S subunit binds to the cap and then scans through the 5□ UTR. The first start codon encountered is at the pORF, leading to assembly of the 80S ribosome and translation initiation. (**B**) When a uORF is present in the 5□ UTR, translation initiation may occur. When the ribosome encounters a termination codon, it may undergo recycling, whereby the 40S and 60S subunits dissociate and the downstream pORF is not translated. (**C**) After translating a uORF, the ribosome is either recycled or the 40S subunit remains associated with the mRNA and scanning continues. If a second uORF is encountered, and translation initiation occurs, then a second ribosome recycling/40S retention and continued scanning event will also occur at the corresponding termination codon. An increase in the frequency of ribosome recycling is associated with a higher degree of pORF translational repression. (**D**) Following uORF translation termination, the 40S subunit must be recharged with the ternary complex (eIF2–GTP–Met-tRNA_i_^Met^) before it is competent to reinitiate translation. If the pORF start codon is encountered during this recharging period it will not be translated, leading to translational repression of the pORF. (**E**) Conversely, uORFs that are positioned far from the pORF may exert minimal repressive effects as the ribosome is permitted to fully recharge after uORF translation termination. (**F**) If a uORF overlaps the pORF start codon and is translated, then the pORF start codon will not be encountered in a translation initiation competent state, leading to pORF translational repression. (**G**) In some cases (such as with homopolymeric acid amino acid-containing artificial uORFs) the ribosome may stall, leading to incomplete/slow translation and a reduced frequency of ribosome recycling or 40S retention and continued scanning events. As a result, fewer translation competent ribosomes encounter the pORF start codon, leading to translational repression.

In cases whereby the 40S subunit continues scanning after a uORF termination codon, there is a refractory period where the ribosome is incompetent to reinitiate translation until it is recharged with the ternary complex (consisting of eIF2–GTP–Met-tRNA_i_^Met^).^74^ If the 40S subunit reaches the pORF start codon without being recharged with the ternary complex, then translation initiation will not occur and the transcript will be translationally repressed with respect to the pORF (**Figure 12D**).^8–10,36,75^ The length of the refractory period has been estimated to be 10-100 nucleotides following a stop codon.^36,75,76^ By contrast, a uORF that is positioned far from the pORF, with sufficient time/distance to become fully recharged and competent to reinitiate, will tend to exhibit negligible, or minimal, translational repressive effects on the downstream pORF (**Figure 12E**), as observed for naturally-occurring cap-proximal uORFs in *UTRN*, *SMN2* (**Figure S4**), and *CUL2* (**Figure 10B**). An interesting implication of this ribosome behavior is the notion that uORFs have the potential to repress one another in multi-uORF contexts. This is exemplified by the well-studied example of *ATF4*,^37,39,40,60^ whereby uORF-2 acts to suppress the activity of uORF-3 (which itself is highly pORF-repressive; **Figure 11D,E**). As such, pORF output in complex 5□ UTRs with multi-uORF configurations is likely to be governed by a combination of initiation probabilities, ribosome recycling rates, and intercistronic distances.

A related concept concerns uORFs which overlap with the pORF, of which there are a substantial number in the human (13,951; 15.3%) and mouse (6,462; 15.1%) transcriptomes (**Figure 1D**) and which are similarly repressive compared to non-pORF-overlapping uORFs (**Figure S9**). In these cases, initiation at the uORF start codon means that the 80S ribosome is elongating and in the process of generating the uORF peptide at the time that the pORF start codon is encountered, meaning that pORF translation cannot initiate (**Figure 12F**).

In some cases, the composition of a uORF peptide may influence its degree of repressive activity. For the native *HOXA11* uORF, the impact of any single amino acid position on uORF activity was found to be minimal (**Figure 5A**). However, artificial homopolymeric uORFs were found to have a pronounced effect on uORF activity, which was strongest for poly-acidic homopolymeric uORFs (**Figure 5B,S11A,S12,7D,8A**). Interestingly, an artificial uORF consisting of repeating glutamic and aspartic acid residues was found to be highly repressive, even when placed in a cap-proximal region (in which the *HOXA11* uORF was found to be non-repressive) (**Figure 8A,9C**) suggestive of a distinct mechanism of action from those described above (**Figure 12A-F**). We propose that these data are explained by a stalling mechanism, whereby translation initiation occurs at the artificial uORF but elongation and/or translation termination are inhibited/delayed. As such, the frequency of both ribosome recycling and 40S subunit continued scanning are reduced. Furthermore, in a stalling model the progression of upstream ribosomes through the transcript is also impeded, resulting in an overall reduction in ribosome flux and a high degree of downstream pORF repression (**Figure 12G**). Ribosome stalling may be a result of differential uORF peptide interactions with side chains in the ribosome nascent peptide channel, or alternatively be driven by the differential rate of peptide bond formation between specific pairs of amino acids.^77–80^ Interestingly, a role for the uORF coding sequence in modulating the speed of uORF translation has been proposed previously in the context of yeast.^81^ Notably, it is possible that other mechanisms may explain these data, for example, if poly-acid uORFs promote very efficient ribosome recycling.

While our machine learning analysis identified amino acids associated with more repressive uORF activity, poly-acidic uORFs were not found to be overrepresented, as might be expected (**Figure 6H,7D**). However, direct analysis of Poly-E uORFs of various lengths did reveal a progressive decline in MRL with length, although the effect sizes were small (**Figure S17**) compared with the experimental data from luciferase reporters (**Figure S11A**). Notably, the Optimus 5-Prime model is trained on polysome profiling data,^45^ whereby MRL is assumed to be a proxy for translation efficiency. If stalling occurs, then ribosomes may remain associated with an mRNA while translation efficiency is low. As such, the Optimus 5-Prime model may underestimate the effects of uORFs which induce ribosome stalling, as we predict for poly-acidic artificial uORFs.

Ribosome stalling at specific amino acid motifs and at sequences rich in proline or basic amino acids (i.e. arginine and lysine) has been reported previously.^82,82–86^ Consistently, we found that poly-arginine and poly-proline synthetic homopolymeric uORFs were highly repressive, although to a lesser extent than the poly-acidic uORFs (**Figure 5B**). Notably, uORFs containing runs of the same repeated amino acid residues were not found in naturally occurring uORFs. Similarly, the random sampling of amino acids in our machine learning datasets also lacked uORFs with extended homopolymeric stretches. It is tempting to speculate that the biophysical properties of amino acid side chains are the key determinant of ‘staller’-like activity, which appears to be the case for poly-acid artificial uORFs (**Figure 5B,S12A,S12B**). However, a link between amino acid biophysical properties and uORF activity is less clear in other cases. For example, we initially observed that homopolymeric artificial uORFs consisting of aromatic hydrophobic sidechains (i.e. phenylalanine and tyrosine) were less-repressive than the native *HOXA11* uORF (**Figure 5B**). However, based on machine learning predictions we observed that poly-tryptophan was more repressive than the WT *HOXA11* uORF (**Figure 6H,7D**). Similarly, poly-tyrosine and poly-serine exhibited opposing effects on uORF activity despite their side-chain similarity (**Figure 7D**). As such, the results reported here with the hyperfunctional artificial uORFs may represent ‘edge cases’, although specific amino acid sequences resulting in dynamic ribosome stalling in response to various stimuli have been described in multiple different organisms.^87–89^

In summary, we propose that repressive uORFs act to inhibit translation via one or more of the following mechanisms; (i) promoting ribosome recycling such that the 40S subunit never reaches the pORF (i.e. ‘recyclers’), (ii) preventing downstream reinitiation events through incomplete ternary complex recharging (i.e. ‘rechargers’), (iii) if an actively-translated uORF overlaps with the pORF then the ribosome will be incompetent to initiate at the pORF start codon (i.e. ‘overlappers’), and (iv) ribosome stalling which delays both recycling and continued 40S subunit scanning (i.e. ‘stallers’).

The data presented herein also point to a hierarchy of importance for uORF properties. Translation initiation is the single most important determinant of uORF functionality as start codon disruption leads to the greatest levels of de-repression (**Figures 2B,3B,4A,5,8A,8D,9C,10B**). Modulating the *HOXA11* uORF Kozak context to optimize it resulted in small increases in repressive activity which did not reach statistical significance at the *P*<0.05 level (**Figure 4A**). Notably, the native *HOXA11* uORF Kozak context is already relatively strong (i.e. an A at the −6 position)^90^, meaning that there may be limited scope to improve the probability of translation initiation at its start codon. Transcriptome-wide analysis showed that strong Kozak context uORFs were found to be more repressive than weak Kozak context uORFs (*D*=0.063, **Figure 4B**), consistent with previous reports,^1,41^ although this result was not statistically significant in our data. The importance of Kozak consensus sequence strength was also indicated in our machine learning analysis, demonstrating that the Optimus 5-Prime algorithm^45^ has learned that this is an important uORF feature. This makes sense intuitively, as the strength of the Kozak context sequence serves to modify the probability of translation initiation at the uORF, with a high probability of initiation associated with a higher degree of repression. According to the model proposed above (**Figure 12**) the importance of Kozak context strength in influencing uORF repressive activity is predicted to depend on the location of the uORF within the 5□ UTR. Specifically, if the translation initiation probability is increased for a pORF-proximal uORF, then both ‘recycler’ and ‘recharger’ inhibition mechanisms are likely to be enhanced. Conversely, for a cap-proximal uORF that terminates >100 nucleotides from the pORF, an increase in Kozak context strength would affect only the ‘recycler’ inhibitory mechanism. Therefore, the influence of Kozak context on uORF repressive activity is predicted to be greater for uORFs that terminate within ∼100 nucleotides of the pORF start codon, according to this model.

Indeed, uORF position was found to be a highly important determinant of uORF activity (**Figure 8,9**), such that uORFs that are repressive in the pORF-proximal position were found to be non-functional in the cap-proximal position (**Figure 8D,S2,9C,10B**). These data suggest that while uORF start codon initiation is essential for uORF-mediated repressive activity, this can be overcome via placement in a cap-proximal 5□ UTR location whereby the uORF can no longer influence initiation at the pORF start codon. Interestingly, these data suggest that, at least in the *BRCA2* and *CUL2* systems we have used, the ‘recharger’ mechanism predominates. (This is because a ‘recycler’ type mechanism should be operative irrespective of 5□ UTR location).

While cap-proximal locations in which uORFs are typically not repressive (likely due to sufficient intercistronic distances to permit complete ternary complex recharging), this effect can be superseded by altering the amino acid content such that ‘staller’ mechanisms are promoted. This appears to be the case for hyperfunctional poly-acidic artificial uORFs which are highly repressive in locations where the native *HOXA11* uORF exhibited negligible repressive activity (**Figure 8A,8D,9C**).

The contribution of each mechanism (i.e. the relative probabilities of ribosome recycling, post-termination continued 40S subunit scanning, or ribosome stalling) is likely to depend on a number of features including uORF sequence composition, length,^74^ and other surrounding features (e.g. 5□ UTR secondary structure).^91^ More detailed mechanistic experiments will be required to determine the contribution of each mechanism to the repressive activity of a given uORF on a case-by-case basis.

We have also investigated other uORF parameters that were found to exhibit minimal influence on *HOXA11* uORF activity and/or in global uORF activity by CDF analysis. In summary, we found that uORF-pORF relative reading frame (**Figure S7**) and rare amino acid composition/availability (**Figure S13**) contributed minimally to uORF-mediated translational repression. Analysis of stop codon usage suggested that UGA is associated with greater uORF repressive activity, although altering the stop codon in the *HOXA11* context had no effect (**Figure S8**). Notably, a UGA stop codon at uORFs has also been reported to be associated with enhanced nonsense mediated decay.^42^

We have utilized luciferase reporter constructs to investigate uORF function, an approach which, although commonly used in uORF research,^1,41^ is an artificial system. The convenience of these assays permits the facile analysis of multiple uORF properties in parallel. Conversely, equivalent analyses in endogenous settings are challenging if a cell model expressing the transcript of interest is unavailable, or if the gene of interest consists of multiple transcripts with differing 5□ UTR configurations and/or uORF complements. In these cases, direct manipulation of uORFs, for example using CRISPR-Cas9, is not possible or may produce ambiguous results.

The interaction between distinct uORF features presents a challenge for studying the influence of a feature in isolation. For example, imposing a strong Kozak consensus context on a uORF start codon inevitably results in the next codon starting with a G nucleotide, thereby limiting the second encoded amino acid to a V, A, D, E, or G. Similarly, for experiments where the location of uORFs is altered, these may result in a shift in both uORF start and stop codons. Conversely, extending the length of a uORF changes not only the length, but also the amino acid composition and potentially the intercistronic distance between uORF and pORF. An important additional consideration is secondary structure, which may be inadvertently altered as a consequence of experimental changes to the sequence of a 5□ UTR of interest.

The CDF analyses contained in this study are necessarily limited to naturally-occurring transcripts for which there is experimental evidence. In contrast to luciferase construct experiments, there are no perfect controls for the CDF analyses, which can only compare populations of existing transcripts with differing properties. Members of these populations may exhibit combinations of properties that can be potentially confounding. For example, the investigation of the role of uORF Kozak context strength on uORF repressive activity may be confounded if some uORFs are located in non-functional 5□ UTR positions, such that changing the uORF Kozak context strength has no influence on the downstream pORF.

Where a uORF property spans a wide range of values (as with uORF length; **Figure 4D**), there is a risk of sample binning being somewhat arbitrary. In this case, our binning strategy was motivated by reflecting the distribution of uORF lengths (**Figure 1C**) while also maintaining similar sample sizes. Indeed, increasing the number of analysis bins necessarily leads to a decrease in statistical power as the total number of transcripts is finite, which constitutes a technical limitation of this approach. This challenge precludes certain analyses that might be informative. For example, it would be interesting to further subset analysis bins for strong and weak Kozak context uORFs according to their relative positions within the 5□ UTR (for which statistical power to detect differences between groups is limited by prohibitively small analysis bin sizes).

Interestingly, we have observed that while aggregated Ribo-Seq can be useful in identifying *bona fide* repressive uORFs (**Figure 3A**), these data can often be misleading. Specifically, Ribo-Seq signal is sometimes observed at uORFs which exert minimal, or negligible, repressive effects on downstream pORFs (e.g. *CUL2*; **Figure 10A**, and *BRD3*; **Figure 11B**, and *ATF4*; **Figure 11D**). These observations may reflect cell-type-specific differences in uORF-ribosome occupancy, that are not captured in the Ribo-Seq datasets used here. Another possibility is that the use of translation inhibitors in the Ribo-Seq protocol (i.e. cycloheximide and harringtonin) may influence the position of ribosomes in a manner that introduces bias in uORF-ribosome occupancy.^92,93^

In summary, we have investigated multiple uORF features and how they influence translational repression activity. We present a model that explains uORF activity through a combination of distinct repressive mechanisms. This work serves as a basis for further investigation of uORF-uORF interactions and the role of other 5□ UTR features in the regulation of pORF translation.

## Methods

### uORF prediction

uORFs were predicted using custom R and Python scripts. The coordinates (i.e. BED files) of all protein-coding transcripts (i.e. those with ‘NM_’ RefSeq IDs) were downloaded using the UCSC Table Browser data retrieval tool^94^ for the human (GRCh38/hg38) and mouse (GRCm38/mm10) genomes. The sequence of all transcripts was retrieved using the getFASTA bedtools (v2.29.2) function.^95^ The 5□ UTR of each construct was searched for upstream canonical start codons (i.e. uATG trinucleotides, uATGs), and the full length transcript was searched for putative stop codons (TAA, TAG, or TGA). The output was limited to in-frame stop codons (using the modulo operator, i.e. %%3 = 0). The minimal possible uORF was defined as 6 nt in length (i.e. methionine-Stop). Sequence logos were generated using WebLogo.^96^ Other outputs from the bioinformatic search included the RNA space co-ordinates (i.e. position along the length of the mature transcript) which enable the determination of the uORF length and whether it spans the translation initiation site of the pORF. The gDNA co-ordinates were calculated such that BED files of uORF predictions could be generated for visualization, and the bigWigAverageOverBed v2 (https://genome.ucsc.edu/goldenpath/help/bigWig.html) program used to calculate the mean phastCons scores for each interval. Visualization of 5□ UTR/uORF configurations together with Ribo-Seq/RNA-Seq data (downloaded from GWIPS-viz)^50^ remapped in RNA space was performed using custom, in-house python scripts.

### Plasmid generation

Plasmids were generated by standard recombinant DNA technology methods or synthesized as a service by Genewiz (Azenta) using custom cloning vectors (**Figure S3**).

### Cell culture

Human embryonic kidney 293T (HEK293T) cells were grown in Dulbecco’s Modified Eagle Medium containing glucose (DMEM, Thermo Fisher Scientific, MA, USA), supplemented with 10% Fetal Bovine Serum (FBS) and 1% Antibiotic-Antimycotic (both Thermo Fisher Scientific). Cells were maintained in a humidified incubator at 37°C with 5% CO_2_. Cell culture stocks were confirmed to be free of *Mycoplasma* contamination by weekly testing. For transfection, 1×10^5^ cells were seeded per well (24 well plate) in 0.5 ml of complete media and treated with 500 ng of plasmid DNA. Transfections were performed using Lipofectamine 2000 (Thermo Fisher Scientific) according to manufacturer’s instructions, and cells harvested 24 hours post transfection.

### Dual luciferase reporter assay

Dual-Glo Luciferase Assays (Promega, WI, USA) were performed 24 hours post transfection to assess the effect of uORF activity on *Renilla* luciferase expression. Firefly luciferase expression was measured as an internal normalization control. Briefly, cells and reagents were equilibrated to room temperature, half of the well volume was removed, and an equal volume of Dual-Glo Reagent was added to each well. Samples were mixed thoroughly by pipetting, and incubated at room temperature for 20 minutes before the lysate was transferred to a Greiner 96 well plate (Merck, NJ, USA). Black plates were used to reduce luminescence cross-talk between wells. Firefly luciferase expression was measured using the CLARIOstar plate reader (BMG Labtech, Aylesbury, UK; at a focal height of 9.0 mm, with no filter), and then quenched by addition of an equal volume of Stop & Glo Reagent (prepared as a 1:100 dilution in Stop & Glo buffer) to the initial culture medium. After 10 minutes at room temperature, *Renilla* luciferase expression was measured using the plate reader (with the same parameters as above). *Renilla* luciferase measurements were normalized to the levels of Firefly luciferase. All luciferase assay experiments were performed at least twice.

### RT-qPCR

Total RNA was extracted using the Maxwell RSC simplyRNA Tissue Kit (Promega) following manufacturer’s instructions. Reverse transcription (random-primed) was performed using the High-Capacity cDNA Reverse Transcription Kit (Applied Biosystems, MA, USA) according to manufacturer’s instructions, with minor modification. Each reaction contained 400 ng of input total RNA in 20 µl total volume was incubated as follows: 10 minutes at 25°C, 30 minutes at 37°C, and 5 minutes at 85°C. qPCR was performed with technical duplicates on a StepOne Plus real-time PCR Thermocycler (Life Technologies, CA, USA) using Power SYBR Green Master Mix (Thermo Fisher Scientific) according to manufacturer’s instructions. Sequences of qPCR primers are listed in **Table S1**. Each PCR reaction consisted of 2 µl of undiluted cDNA in a 20 µl total reaction volume. Cycling conditions were as follows: initial denaturation for 10 minutes at 95°C, followed by 40 cycles of 15 seconds at 95°C and 1 minute at 60°C. Reaction specificity was confirmed by post-run melt curve analysis. Relative quantification was performed using the Pfaffl method.^97^

### uORF peptide detection

Levels of HiBiT-tagged uORF-encoded micropeptides were measured 48 hours post transfection by nanoluciferase biocomplementation assay using the Nano-Glo HiBiT Lytic Detection System (Promega). Plates and buffers were first equilibrated to room temperature before half of the well volume was removed, and an equal volume of Nano-Glo HiBiT Lytic Reagent (1:100 of LgBiT Protein, and 1:50 of Nano-Glo HiBiT Lytic Substrate in Lytic Buffer) was added to the remaining culture medium. Samples were mixed thoroughly by pipetting and incubated for 10 minutes at room temperature. NanoLuc expression was measured in triplicate, using the CLARIOstar plate reader (at a focal height of 9.0 mm, and an integration time of 1 second). Background expression from untreated cells was subtracted from the values obtained.

For western blotting, cell lysates were incubated at 95□°C for 5□minutes in 1× Laemmli sample buffer. Samples were and separated by SDS polyacrylamide gel electrophoresis using 4-12% gradient polyacrylamide pre-cast gels (Thermo Fisher Scientific). Proteins were electrotransferred onto a polyvinylidene fuoride membrane by electroblotting and immunoblotted using an anti-FLAG antibody (F1804; Sigma-Aldrich) using standard procedures.

For both nanoluciferase biocomplementation assay and anti-FLAG western blot, samples were analysed in both the presence and absence of the 26S proteasome inhibitor MG132 (10 μM; Sigma Aldrich).

### *In vitro* transcription/cell-free translation assay

Plasmid constructs were amplified by PCR to generate templates for *in vitro* transcription (IVT) whereby a T7 promoter sequence was introduced at the 5□ terminus. Synthetic mRNAs were synthesized using the HiScribe T7 ARCA (New England Biolabs) IVT kit according to manufacturer’s instructions with 2 µg of input template DNA. Reactions were thoroughly mixed and briefly centrifuged before being incubated at 37°C for 60 minutes. Subsequently, 2 U of DNase I (New England Biolabs) was then added to each reaction and incubated at 37°C for 15 minutes. Poly(A)-tailing was subsequently performed using 5 U of Poly(A) Polymerase in Poly(A) Polymerase Reaction Buffer (both New England Biolabs) and incubated at 37°C for 45 minutes. Capped and tailed synthetic mRNAs were isolated using the MegaClear Clean-Up kit (Thermo Fisher Scientific) following the manufacturer’s instructions.

Cell-free translation (CFT) was performed using rabbit reticulocyte lysate (RRL; Promega). Briefly, reactions consisted of 70% (v/v) RRL, 20 µM Amino Acid Mixture, 0.8 U/µ□ RNase Inhibitor, and 20 ng/µ□ of each luciferase mRNA (i.e. either control Firefly luciferase or experimental *Renilla* luciferase). Samples were immediately incubated at 30°C for a total of 90 minutes, with agitation (300 rpm). Following CFT, samples were analysed by dual luciferase assay as described above.

### Bioinformatics

Cumulative distribution function (CDF) analyses were performed to assess the relationship between various uORF features and global protein expression. These analyses utilized matched proteomics and transcriptomics data from 29 healthy human tissues as described by Wang *et al*.^44^ Briefly, transcripts were first binned according to uORF features as indicated, and the CDFs of the corresponding protein expression values for each bin were used to evaluate the importance of each feature on uORF activity. Given that the protein data lack isoform resolution, the median protein expression was used as representative value for protein expression for each gene. For transcriptomic analysis, data were filtered to include only transcripts for which protein expression data were available. For the majority of the analyses, we focused on transcripts that contained only a single uORF (as indicated), as more complex 5□ UTR/uORF architectures can potentially be confounding. Where genes consisted of transcripts which could be assigned to more than one bin (e.g. as a consequence of alternative splicing or alternative promotor usage) they were excluded from the analysis.

Machine learning analysis was performed using a published model, Optimus 5-Prime.^45^ Generation of simulation sets and implementation of the model predictions were performed using custom Python scripts.

The metagene analysis was performed using deepTools (v.3.5.1). A BED file of the human transcriptome 5□ UTRs was generated in the UCSC table browser. The computeMatrix software was performed on the ribosome profile and ribosome initiations aggregated files, using the 5□ UTR BED file. computeMatrix was run with the scale-regions parameter, and the 5□ UTR regions were binned to 100 bins. plotProfile analysis was used to plot the aggregate the matrices and generate figures.

### Statistics

Statistical analysis was performed using GraphPad Prism 10.1.2 (GraphPad Software Inc., San Diego, California, USA). Two sample comparisons were analysed using a Student’s *t*-test. Comparisons between more than two groups were assessed by one-way analysis of variance (ANOVA) and Bonferroni *post hoc* test. Analyses were performed on log_2_-transformed expression ratios. Two-sample Kolmogorov-Smirnov tests and χ^2^ tests were performed in Python using the *scipy.stats.ks_2samp* and *scipy.stats.chisquare* functions from the SciPy library.^98^ Sequence logos were generated in Python using the *logomaker*library.^99^

## Acknowledgements

This work was supported by grants from Great Ormond Street Hospital Sparks Fund/Dravet Syndrome UK (V4121) and UK MRC (TransNAT) (MR/X008029/1) (awarded to MJAW and TCR), the Oxford University Press John Fell Fund and Medical Life Sciences Translational Fund (awarded to TCR). BH is supported by a doctoral studentship from the Clarendon Fund and a grant from the Medical Life Sciences Translational Fund (MLSTF) awarded to TCR. KC was supported by doctoral studentship from the Clarendon Fund in partnership with the Medical Research Council (MRC), and the Juel-Jenson Scholarship from St Cross College, Oxford.

## Author Contributions

TCR, BH, NS, SEA, and MJAW conceived the study. TCR and MJAW supervised the work. BH, NS, NF, HJF, AJL, RA, NH, KC, MD, YJ performed experimentation. TCR wrote the first draft of the manuscript. All authors contributed to the final version of the manuscript.

## Declaration of Interests

TCR, MJAW and BH have filed a patents related to a uORF-disrupting technologies. TCR, MJAW, NS, and BH are founders and shareholders in Orfonyx Bio Ltd, a biotechnology spin-out company that aims to utilize uORF-targeting technologies for therapeutics development. NS is an employee of Orfonyx Bio. TCR and MJAW are consultants for Orfonyx Bio.

## Data Availability Statement

All data are included in the manuscript. Raw data are available on request.

