## Supplementary Information for "Toward a model of uORF-mediated translational control: An integrated bioinformatic and experimental approach"

| | FWD | REV |
|---|---|---|
| **qRLuc** | GTAACGCTGCCTCCAGCTAC | CCAAGCGGTGAGGTACTTGT |
| **qFLuc** | ACTCTAAGACCGACTACCAGG | GTAGACCCAGAGCTGTTCATG |

**Table S1**

**List of primers used in this study.**

All primer sequences are shown in the 5′ to 3′ orientation.

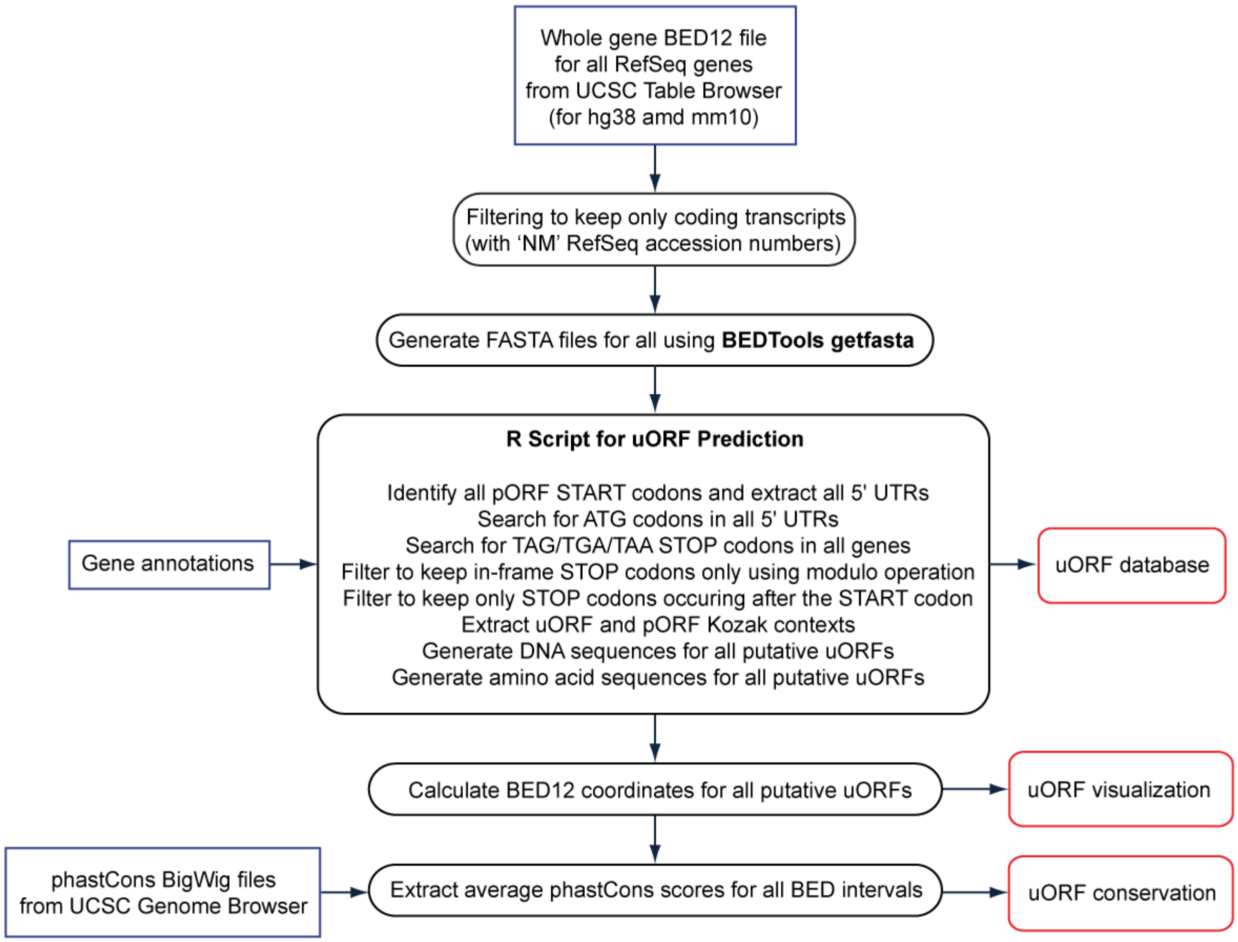


**Figure S1**

**Prediction of uORFs in human and mouse transcripts.**

Schematic of bioinformatics pipeline for uORF prediction. Bioinformatics tools are highlighted in bold. Input data and output resources are highlighted in blue and red boxes respectively.

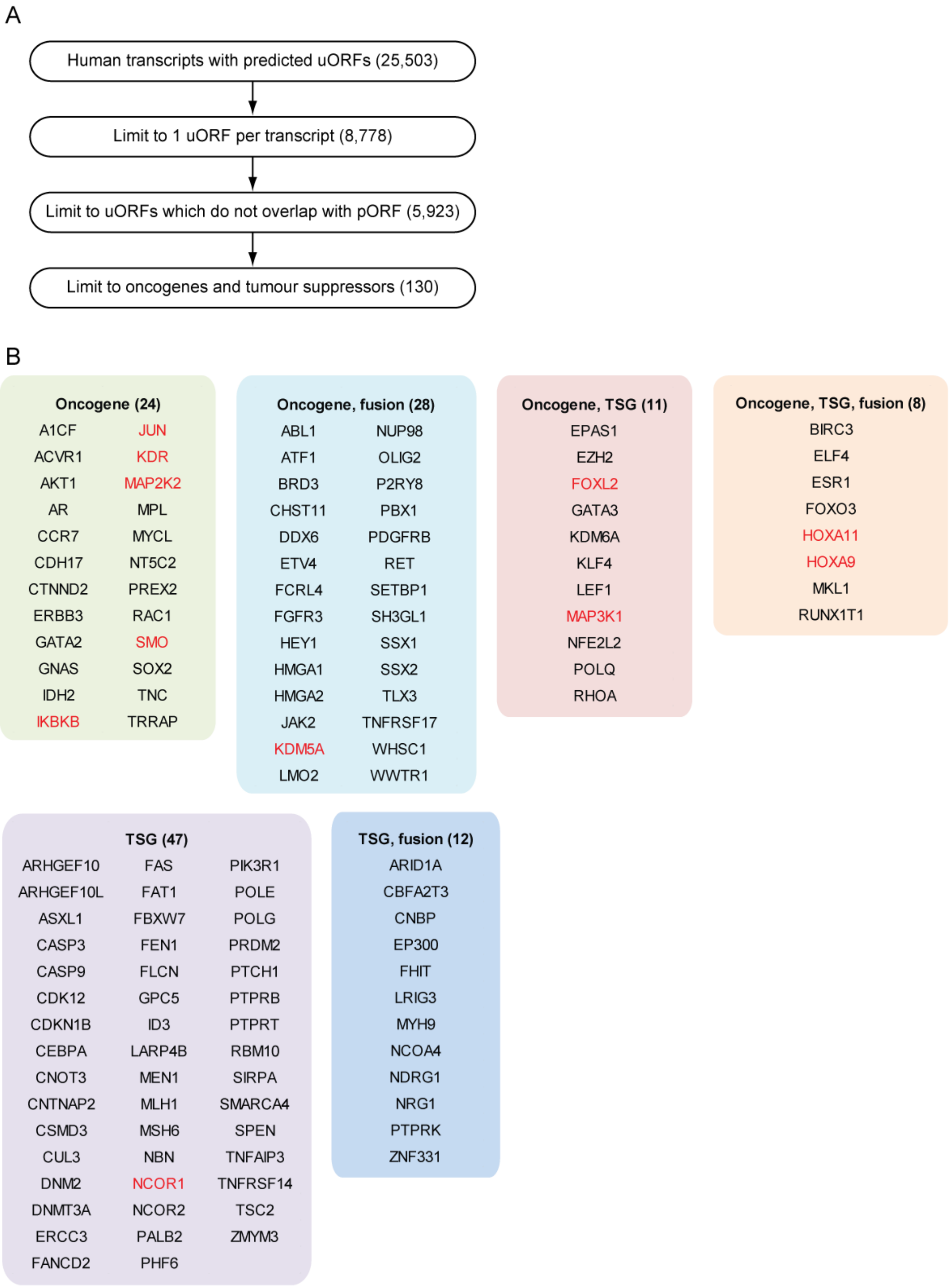
A
Human transcripts with predicted uORFs (25,503)
Limit to 1 uORF per transcript (8,778)
Limit to uORFs which do not overlap with pORF (5,923)
Limit to oncogenes and tumour suppressors (130)
B
Oncogene (24)
A1CF JUN
ACVR1 KDR
AKT1 MAP2K2
AR MPL
CCR7 MYCL
CDH17 NT5C2
CTNND2 PREX2
ERBB3 RAC1
GATA2 SMO
GNAS SOX2
IDH2 TNC
IKBKB TRRAP
Oncogene, fusion (28)
ABL1 NUP98
ATF1 OLIG2
BRD3 P2RY8
CHST11 PBX1
DDX6 PDGFRB
ETV4 RET
FCRL4 SETBP1
FGFR3 SH3GL1
HEY1 SSX1
HMGA1 SSX2
HMGA2 TLX3
JAK2 TNFRSF17
KDM5A WHSC1
LMO2 WWTR1
Oncogene, TSG (11)
EPAS1
EZH2
FOXL2
GATA3
KDM6A
KLF4
LEF1
MAP3K1
NFE2L2
POLQ
RHOA
Oncogene, TSG, fusion (8)
BIRC3
ELF4
ESR1
FOXO3
HOXA11
HOXA9
MKL1
RUNX1T1
TSG (47)
ARHGEF10 FAS PIK3R1
ARHGEF10L FAT1 POLE
ASXL1 FBXW7 POLG
CASP3 FEN1 PRDM2
CASP9 FLCN PTCH1
CDK12 GPC5 PTPRB
CDKN1B ID3 PTPRT
CEBPA LARP4B RBM10
CNOT3 MEN1 SIRPA
CNTNAP2 MLH1 SMARCA4
CSMD3 MSH6 SPEN
CUL3 NBN TNFAIP3
DNM2 NCOR1 TNFRSF14
DNMT3A NCOR2 TSC2
ERCC3 PALB2 ZMYM3
FANCD2 PHF6
TSG, fusion (12)
ARID1A
CBFA2T3
CNBP
EP300
FHIT
LRIG3
MYH9
NCOA4
NDRG1
NRG1
PTPRK
ZNF331

**Figure S2**

**Selection of candidate uORFs for experimental validation.**

(**A**) Schema illustrating the filtering process for validation candidate selection. Human transcripts containing predicted uORFs were filtered to keep only transcripts with a single uORF that did not overlap with the pORF. This list was then intersected with known oncogenes and tumor suppressor genes (TSGs) from the COSMIC database. The number of transcripts at each filtering step are indicated in parentheses. (**B**) Gene names for the 130 filtered candidates are shown separated by category. Potential candidates for experimental validation were manually inspected, and those selected are highlighted in red.

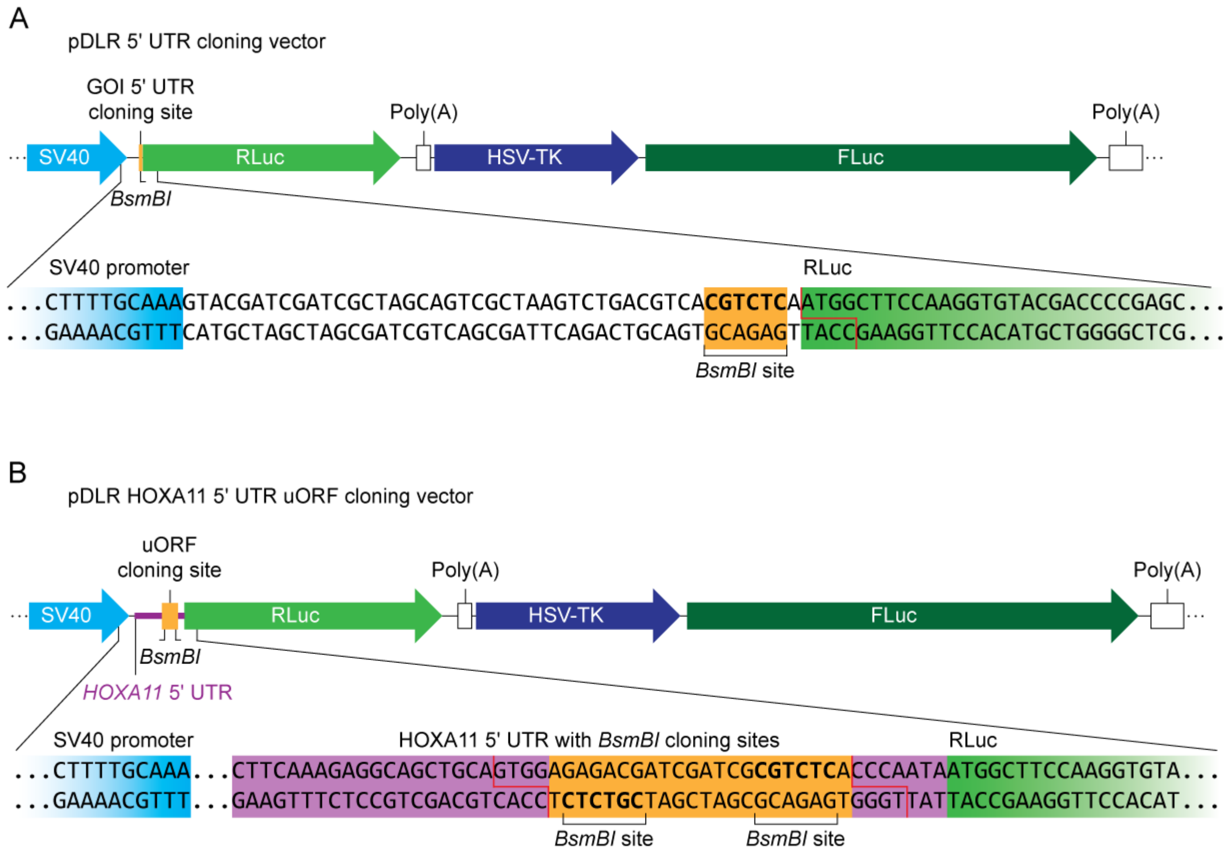


**Figure S3**

**Schematics of in-house cloning vectors used in this study.**

(**A**) pDLR 5′ UTR cloning vector. This vector contains two independent luciferase transgene cassettes: *Renilla* luciferase (RLuc) and firefly luciferase (FLuc). A 5′ UTR for a gene of interest (GOI) can be inserted immediately upstream of the RLuc transgene by restriction cloning at a *BsmBI* site. (**B**) pDLR HOXA11 5′ UTR uORF cloning vector. This variant vector contains the HOXA11 5′ UTR cloned upstream of the RLuc transgene. The HOXA11 uORF and Kozak context are replaced by a Golden Gate cloning site consisting of opposing *BsmBI* restriction sites. Cloning into this site allows for facile testing of various mutant *HOXA11* uORF sequences which can be seamlessly inserted into the *HOXA11* 5′ UTR.

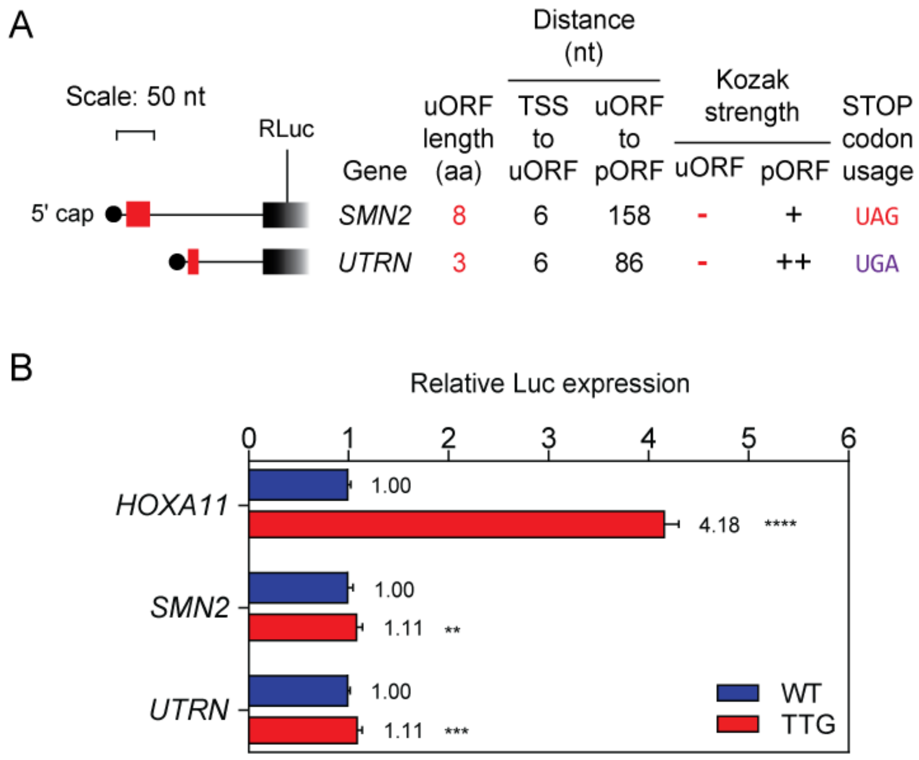


**Figure S4**

**Analysis of non-functional uORFs.**

(**A**) Features of the 5′ UTRs and predicted uORFs for survivor of motor neuron 2 (*SMN2*) and utrophin (*UTRN*). uORFs are depicted by red boxes. Each 5′ UTR was cloned upstream of the RLuc transgene of a dual-luciferase reporter vector (pDLR). uORF-disrupted (TTG) mutants were generated for each 5′ UTR. The *HOXA11* 5′ UTR was analysed in parallel as a control. (**B**) HEK293T cells were transfected with each pDLR construct and luciferase activity measured 24 hours post transfection. Values of the WT and TTG version of each uORF are mean+SEM ($n$=4), and were scaled such that the mean of the WT control group was returned to a value of 1. Statistical significance between each WT and TTG construct was determined by Student's $t$-test. **$P$<0.01, ***$P$<0.001, ****$P$<0.0001. TSS, transcription start site.

**KS-test results for all proteins separated by tissue (presence of uORF vs absence of uORF)**

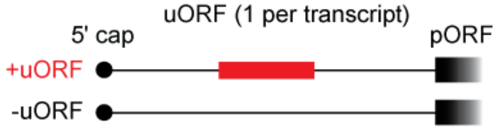


Aggregated proteomics data

| | *D*-statistic | *P*-value |
|---|---|---|
| **Adrenal gland** | 0.140 | 4.61E-38 |
| **Appendix** | 0.132 | 1.27E-30 |
| **Brain** | 0.115 | 7.00E-26 |
| **Colon** | 0.133 | 5.00E-34 |
| **Duodenum** | 0.153 | 2.33E-46 |
| **Endometrium** | 0.141 | 8.89E-38 |
| **Esophagus** | 0.143 | 7.00E-38 |
| **Fallopian tube** | 0.138 | 8.35E-39 |
| **Fat** | 0.142 | 1.42E-34 |
| **Gallbladder** | 0.140 | 3.13E-36 |
| **Heart** | 0.129 | 1.13E-29 |
| **Kidney** | 0.135 | 4.84E-33 |
| **Liver** | 0.125 | 6.07E-29 |
| **Lung** | 0.140 | 1.03E-40 |
| **Lymph node** | 0.127 | 1.72E-29 |
| **Ovary** | 0.143 | 4.30E-37 |
| **Pancreas** | 0.143 | 2.09E-35 |
| **Placenta** | 0.142 | 5.35E-38 |
| **Prostate** | 0.144 | 3.14E-38 |
| **Rectum** | 0.137 | 4.07E-35 |
| **Salivary gland** | 0.138 | 3.91E-35 |
| **Small intestine** | 0.149 | 1.37E-44 |
| **Smooth muscle** | 0.127 | 1.68E-29 |
| **Spleen** | 0.131 | 4.37E-30 |
| **Stomach** | 0.151 | 5.04E-44 |
| **Testis** | 0.132 | 3.94E-34 |
| **Thyroid** | 0.138 | 7.20E-35 |
| **Tonsil** | 0.141 | 7.23E-37 |
| **Urinary bladder** | 0.136 | 2.16E-33 |

**Figure S5**

**Tissue-wide effect of uORFs on protein of the downstream pORF.**

Publicly available matched proteomics and RNA-Seq data from 29 healthy human tissues[44] were analysed and cumulative distributions plotted for protein expression for transcripts containing one or more predicted uORFs versus transcripts containing no uORFs. Statistical significance between distributions was tested by Kolmogorov-Smirnov test and *D*-statistic and *P*-values tabulated. Results were broadly similar between tissues.

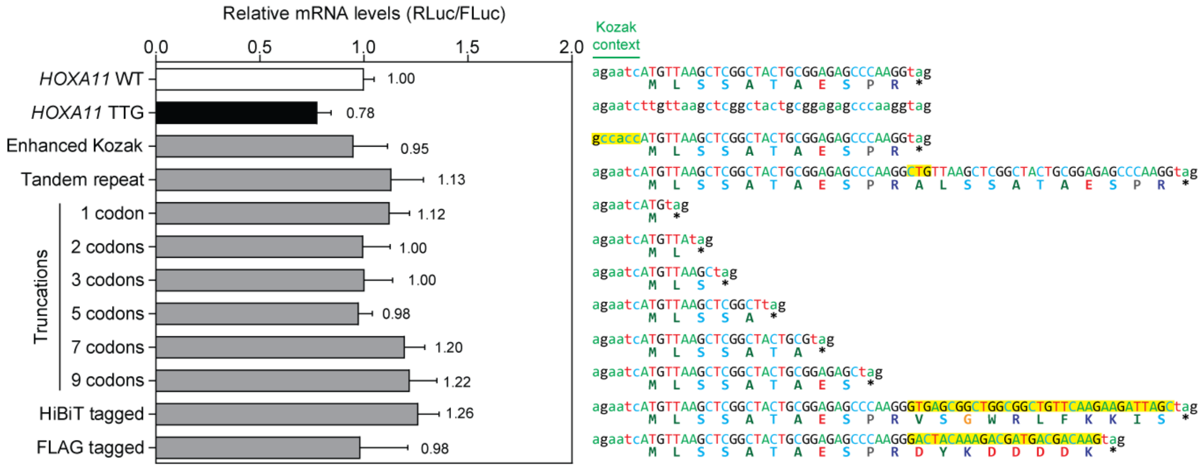


**Figure S6**

**Effect of uORF length and Kozak context sequence on transcript levels.**

HEK293T cells were transfected with pDLR constructs as described in **Figure 3** and relative luciferase transcript levels were determined by RT-qPCR. RLuc expression was normalized to Fluc transcript levels and relative luciferase transcript levels determined 24 hours post transfection. Key mutated bases are highlighted in yellow. Values are mean+SEM, (*n*=3), and were scaled such that the mean of the WT control group was returned to a value of 1. Statistical significance was assessed by one-way ANOVA and Bonferroni *post hoc* test.

A

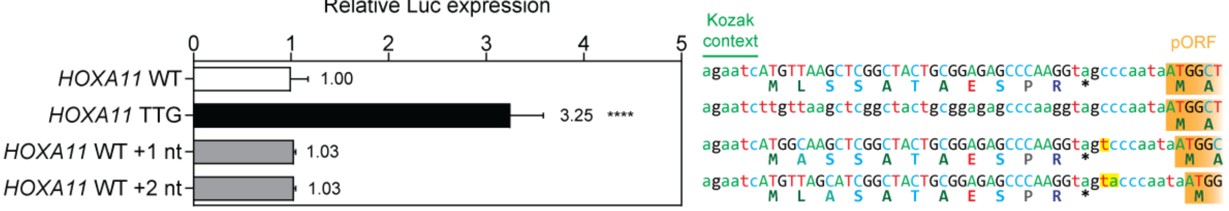


B

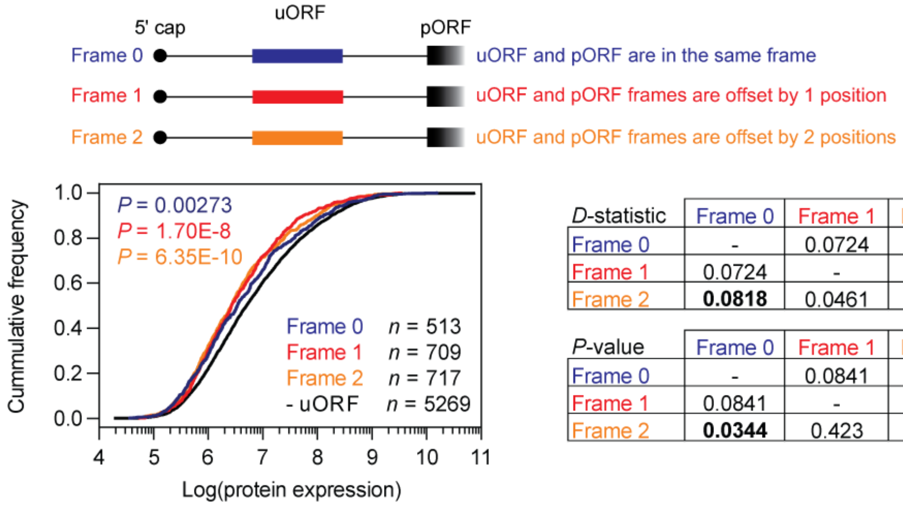


| *D*-statistic | Frame 0 | Frame 1 | Frame 2 | - uORF |
|---|---|---|---|---|
| Frame 0 | - | 0.0724 | **0.0818** | **0.0836** |
| Frame 1 | 0.0724 | - | 0.0461 | **0.121** |
| Frame 2 | **0.0818** | 0.0461 | - | **0.131** |

| *P*-value | Frame 0 | Frame 1 | Frame 2 | - uORF |
|---|---|---|---|---|
| Frame 0 | - | 0.0841 | **0.0344** | **0.00273** |
| Frame 1 | 0.0841 | - | 0.423 | **1.70E-08** |
| Frame 2 | **0.0344** | 0.423 | - | **6.35E-10** |

**Figure S7**

**Analysis of the uORF relative reading frame on translation repression activity.**

HEK293T cells were transfected with various pDLR *HOXA11* 5′ UTR constructs and luciferase activity determined 24 hours post transfection. *HOXA11* WT and a uORF-disrupted (TTG) mutant were included as controls. (**A**) Variants of the *HOXA11* uORF were tested where the difference in reading frame between the uORF and pORF was altered by addition of either 1 or 2 nucleotides. Inserted bases are highlighted in yellow and the start of the pORF is indicated with orange boxes. Values are mean+SEM ($n$=4), and were scaled such that the mean of the WT control group was returned to a value of 1. Statistical significance was determined by one-way ANOVA with Bonferroni *post hoc* test, ****$P$<0.0001. (**B**) Aggregated proteomics data from 29 healthy human tissues were binned according to the relative uORF reading frame (frame 0 is the same frame as the pORF). The numbers of proteins in each bin are indicated. Statistical differences between distributions were assessed by Kolmogorov-Smirnov test with $D$-statistics and $P$-values tabulated.

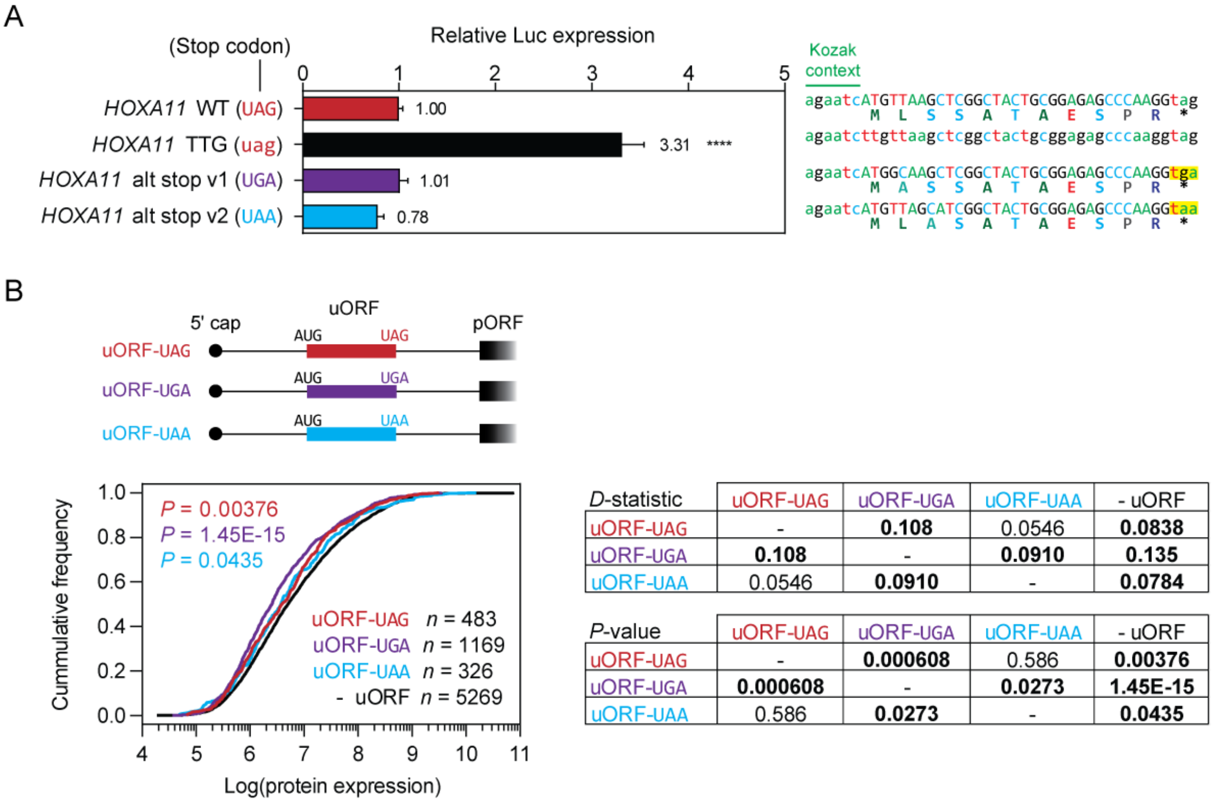


| *D*-statistic | uORF-UAG | uORF-UGA | uORF-UAA | - uORF |
|---|---|---|---|---|
| uORF-UAG | - | **0.108** | 0.0546 | **0.0838** |
| uORF-UGA | **0.108** | - | **0.0910** | **0.135** |
| uORF-UAA | 0.0546 | **0.0910** | - | **0.0784** |

| *P*-value | uORF-UAG | uORF-UGA | uORF-UAA | - uORF |
|---|---|---|---|---|
| uORF-UAG | - | **0.000608** | 0.586 | **0.00376** |
| uORF-UGA | **0.000608** | - | **0.0273** | **1.45E-15** |
| uORF-UAA | 0.586 | **0.0273** | - | **0.0435** |

**Figure S8**

**Analysis of the influence of uORF stop codon usage on translation repression activity.**

HEK293T cells were transfected with various pDLR *HOXA11* 5′ UTR constructs and luciferase activity determined 24 hours post transfection. *HOXA11* WT and a uORF-disrupted (TTG) mutant were included as controls. (**A**) Variants of the *HOXA11* uORF were tested where the stop codon was mutated to TAA or TGA. Key changed codons are highlighted in yellow. Values are mean+SEM ($n$=4), and were scaled such that the mean of the WT control group was returned to a value of 1. Statistical significance was determined by one-way ANOVA with Bonferroni *post hoc* test. ****$P$<0.0001. (**B**) Aggregated proteomics data from 29 healthy human tissues were binned according to the uORF stop codon usage and data presented as cumulative distribution function plots. The numbers of proteins in each bin are indicated. Statistical differences between distributions were assessed by Kolmogorov-Smirnov test with $D$-statistics and $P$-values tabulated.

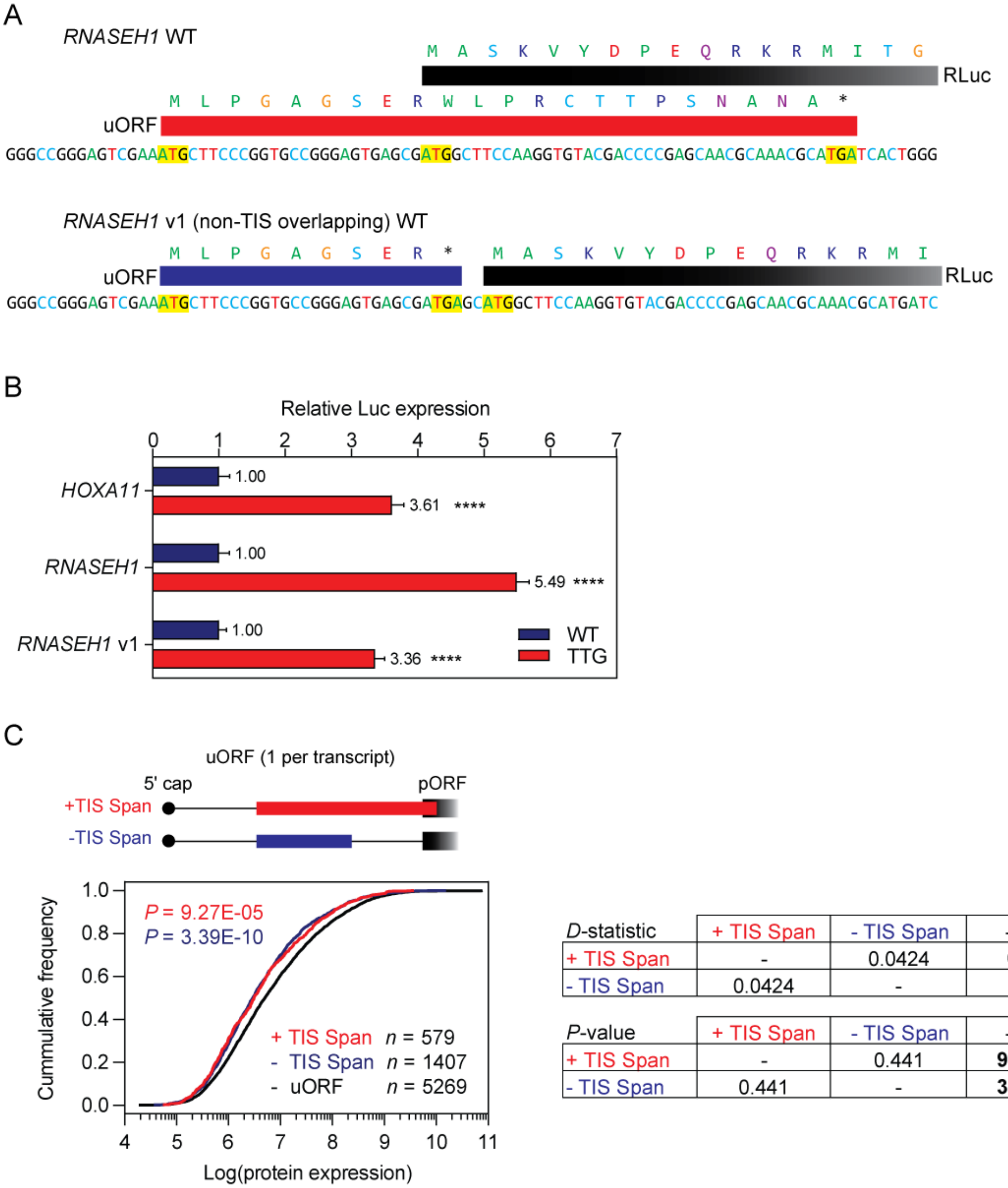


| *D*-statistic | + TIS Span | - TIS Span | - uORF |
|---|---|---|---|
| + TIS Span | - | 0.0424 | **0.0974** |
| - TIS Span | 0.0424 | - | **0.100** |

| *P*-value | + TIS Span | - TIS Span | - uORF |
|---|---|---|---|
| + TIS Span | - | 0.441 | **9.27E-05** |
| - TIS Span | 0.441 | - | **3.39E-10** |

**Figure S9**

**Effect of uORF spanning the translation initiation site on uORF activity.**

(**A**) Design of pDLR constructs: WT *RNASEH1* uORF (TIS overlapping) and *RNASEH1* v1 uORF (non-TIS overlapping). (**B**) HEK293T cells were transfected with the WT and v1 *RNASEH1* constructs and luciferase activity assayed 24 hours post transfection. Values are mean+SEM ($n$=4), and were scaled such that the mean of the WT control group was returned to a value of 1. Statistical significance was determined by Students $t$-test, ****$P$<0.0001. (**C**) Aggregated proteomics data from 29 healthy human tissues were binned according to whether or not a predicted uORF spans the TIS (i.e. the pORF) and data presented as cumulative distribution function plots. The numbers of proteins in each bin are indicated. Statistical differences between distributions were assessed by Kolmogorov-Smirnov test with $D$-statistics and $P$-values tabulated.

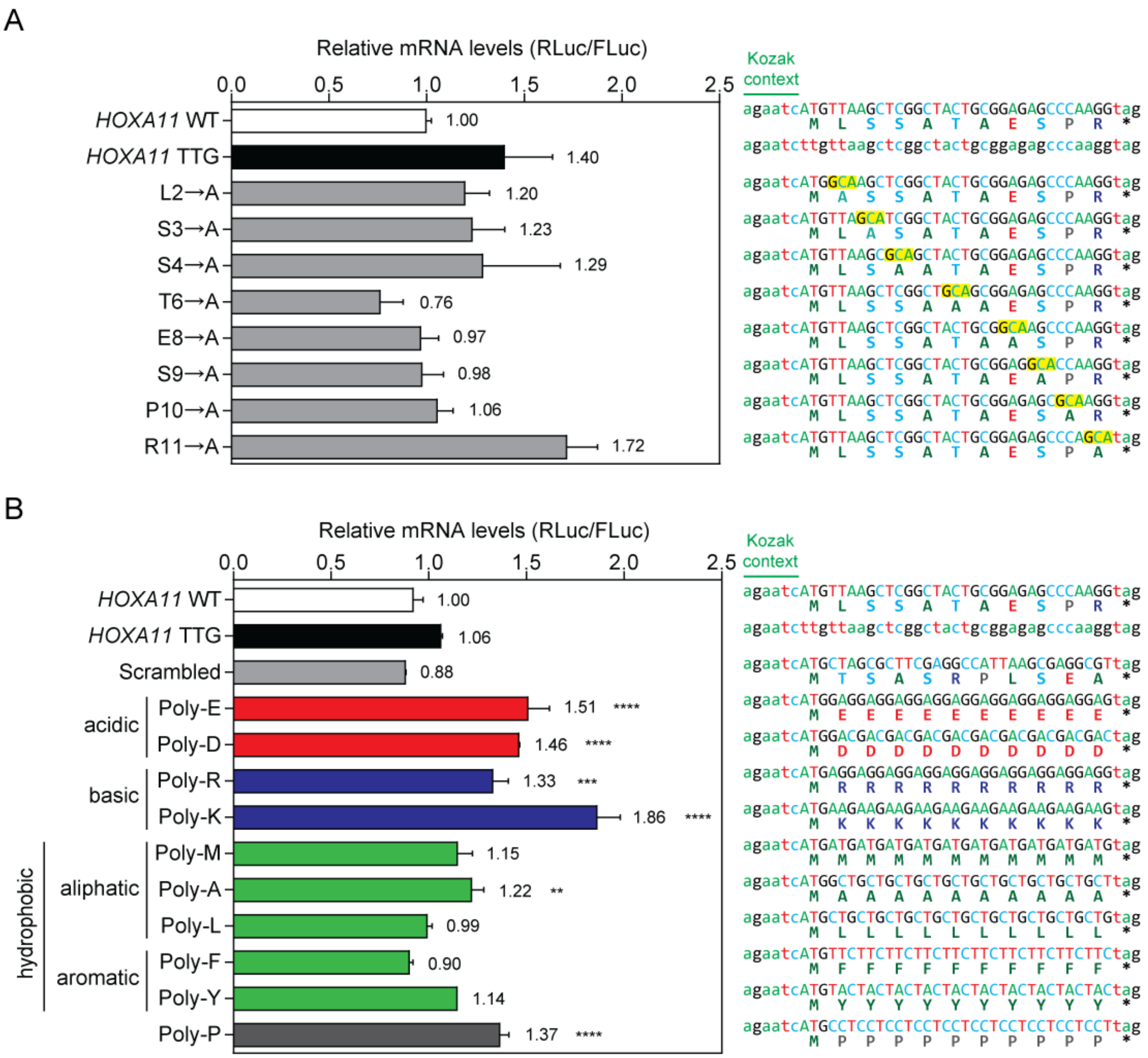
A
Relative mRNA levels (RLuc/FLuc)
0.0
0.5
1.0
1.5
2.0
2.5
Kozak context
HOXA11 WT
1.00
agaatcATGTTAAGCTCGGCTACTGCGGAGAGCCCAAGGtag
M L S S A T A E S P R *
HOXA11 TTG
1.40
agaatcttgttaagctcggctactgcggagagcccaaggtag
L2→A
1.20
agaatcATGGCAAGCTCGGCTACTGCGGAGAGCCCAAGGtag
M A S S A T A E S P R *
S3→A
1.23
agaatcATGTTAGCATCGGCTACTGCGGAGAGCCCAAGGtag
M L A S A T A E S P R *
S4→A
1.29
agaatcATGTTAAGCGCAGCTACTGCGGAGAGCCCAAGGtag
M L S A A T A E S P R *
T6→A
0.76
agaatcATGTTAAGCTCGGCTGCAGCGGAGAGCCCAAGGtag
M L S S A A A E S P R *
E8→A
0.97
agaatcATGTTAAGCTCGGCTACTGCGGCAAGCCCAAGGtag
M L S S A T A A S P R *
S9→A
0.98
agaatcATGTTAAGCTCGGCTACTGCGGAGGCACCAAGGtag
M L S S A T A E A P R *
P10→A
1.06
agaatcATGTTAAGCTCGGCTACTGCGGAGAGCGCAAGGtag
M L S S A T A E S A R *
R11→A
1.72
agaatcATGTTAAGCTCGGCTACTGCGGAGAGCCCAGCAtag
M L S S A T A E S P A *
B
Relative mRNA levels (RLuc/FLuc)
0.0
0.5
1.0
1.5
2.0
2.5
Kozak context
HOXA11 WT
1.00
agaatcATGTTAAGCTCGGCTACTGCGGAGAGCCCAAGGtag
M L S S A T A E S P R *
HOXA11 TTG
1.06
agaatcttgttaagctcggctactgcggagagcccaaggtag
Scrambled
0.88
agaatcATGCTAGCGCTTCGAGGCCATTAAGCGAGGCGTtag
M T S A S R P L S E A *
acidic
Poly-E
1.51 ****
agaatcATGGAGGAGGAGGAGGAGGAGGAGGAGGAGGAGtag
M E E E E E E E E E E *
Poly-D
1.46 ****
agaatcATGGACGACGACGACGACGACGACGACGACGACtag
M D D D D D D D D D D *
basic
Poly-R
1.33 ***
agaatcATGAGGAGGAGGAGGAGGAGGAGGAGGAGGAGGtag
M R R R R R R R R R R *
Poly-K
1.86 ****
agaatcATGAAGAAGAAGAAGAAGAAGAAGAAGAAGAAGtag
M K K K K K K K K K K *
hydrophobic
aliphatic
Poly-M
1.15
agaatcATGATGATGATGATGATGATGATGATGATGATGtag
M M M M M M M M M M M *
Poly-A
1.22 **
agaatcATGGCTGCTGCTGCTGCTGCTGCTGCTGCTGCTtag
M A A A A A A A A A A *
Poly-L
0.99
agaatcATGCTGCTGCTGCTGCTGCTGCTGCTGCTGCTGtag
M L L L L L L L L L L *
aromatic
Poly-F
0.90
agaatcATGTTCTTCTTCTTCTTCTTCTTCTTCTTCTTCtag
M F F F F F F F F F F *
Poly-Y
1.14
agaatcATGTACTACTACTACTACTACTACTACTACTACtag
M Y Y Y Y Y Y Y Y Y Y *
Poly-P
1.37 ****
agaatcATGCCTCCTCCTCCTCCTCCTCCTCCTCCTCCTtag
M P P P P P P P P P P *

**Figure S10**

**Effect of uORF amino acid composition on transcript levels.**

HEK293T cells were transfected with pDLR constructs related to (**A**) single amino acid alanine substitutions, and (**B**) homoploymeric amino acid artificial uORFs as described in **Figure 5** and relative luciferase transcript levels determined by RT-qPCR 24 hours post transfection. RLuc expression was normalized to Fluc transcript levels. Key mutated bases are highlighted in yellow. Values are mean+SEM, ($n$=3), and were scaled such that the mean of the WT control group was returned to a value of 1. Statistical significance was assessed by one-way ANOVA and Bonferroni *post hoc* test. **$P$<0.01, ***$P$<0.001, ****$P$<0.0001.

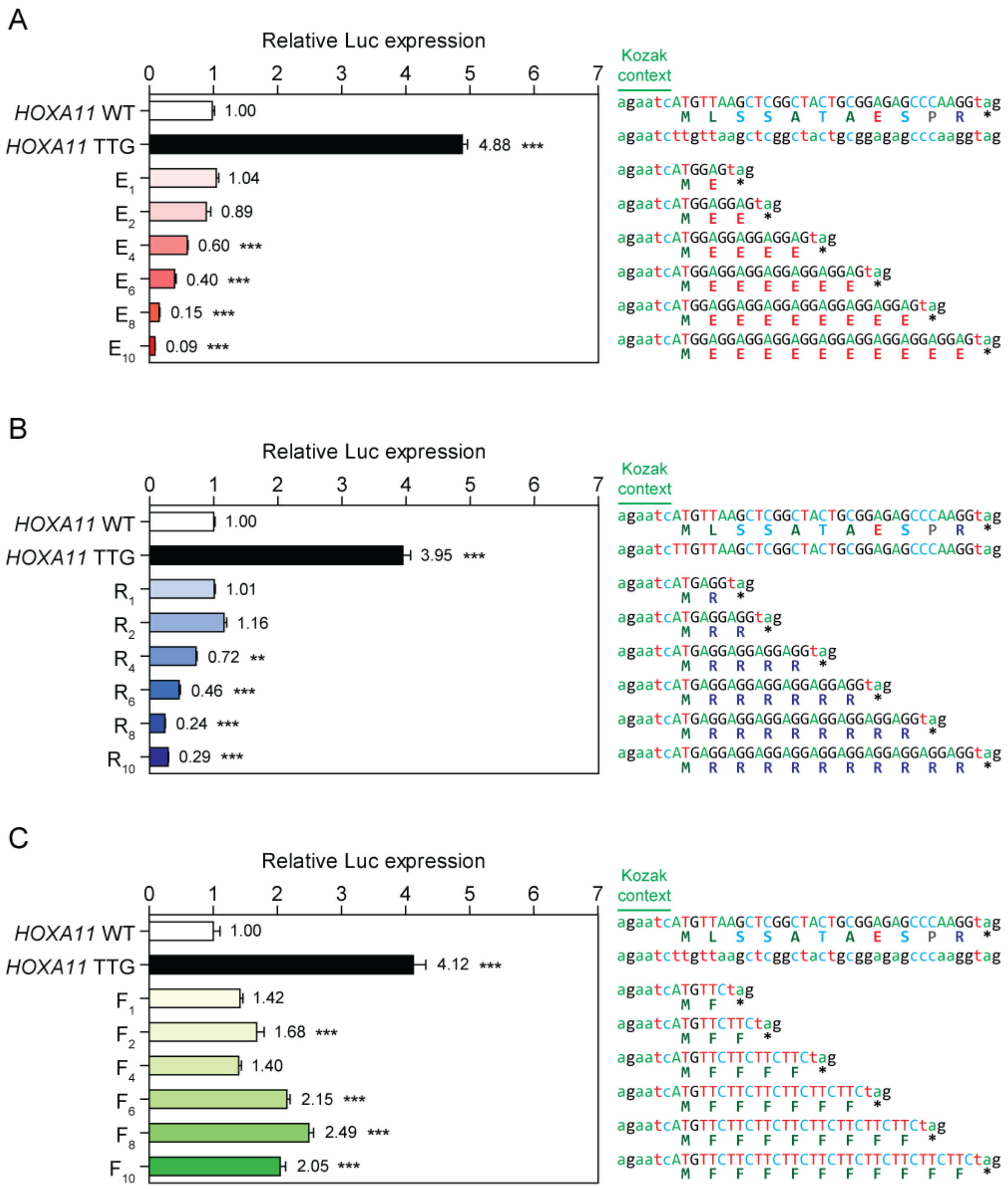

A
Relative Luc expression
0 1 2 3 4 5 6 7
HOXA11 WT 1.00
HOXA11 TTG 4.88 ***
E1 1.04
E2 0.89
E4 0.60 ***
E6 0.40 ***
E8 0.15 ***
E10 0.09 ***
Kozak context
agaatcATGTTAAGCTCGGCTACTGCGGAGAGCCCAAGGtag
M L S S A T A E S P R *
agaatcttgttaagctcggctactgcggagagcccaaggtag
agaatcATGGAGtag
M E *
agaatcATGGAGGAGtag
M E E *
agaatcATGGAGGAGGAGGAGtag
M E E E E *
agaatcATGGAGGAGGAGGAGGAGGAGtag
M E E E E E E *
agaatcATGGAGGAGGAGGAGGAGGAGGAGGAGtag
M E E E E E E E E *
agaatcATGGAGGAGGAGGAGGAGGAGGAGGAGGAGGAGtag
M E E E E E E E E E E *
B
Relative Luc expression
0 1 2 3 4 5 6 7
HOXA11 WT 1.00
HOXA11 TTG 3.95 ***
R1 1.01
R2 1.16
R4 0.72 **
R6 0.46 ***
R8 0.24 ***
R10 0.29 ***
Kozak context
agaatcATGTTAAGCTCGGCTACTGCGGAGAGCCCAAGGtag
M L S S A T A E S P R *
agaatcTTGTTAAGCTCGGCTACTGCGGAGAGCCCAAGGtag
agaatcATGAGGtag
M R *
agaatcATGAGGAGGtag
M R R *
agaatcATGAGGAGGAGGAGGtag
M R R R R *
agaatcATGAGGAGGAGGAGGAGGAGGtag
M R R R R R R *
agaatcATGAGGAGGAGGAGGAGGAGGAGGAGGtag
M R R R R R R R R *
agaatcATGAGGAGGAGGAGGAGGAGGAGGAGGAGGAGGtag
M R R R R R R R R R R *
C
Relative Luc expression
0 1 2 3 4 5 6 7
HOXA11 WT 1.00
HOXA11 TTG 4.12 ***
F1 1.42
F2 1.68 ***
F4 1.40
F6 2.15 ***
F8 2.49 ***
F10 2.05 ***
Kozak context
agaatcATGTTAAGCTCGGCTACTGCGGAGAGCCCAAGGtag
M L S S A T A E S P R *
agaatcttgttaagctcggctactgcggagagcccaaggtag
agaatcATGTTCtag
M F *
agaatcATGTTCTTCtag
M F F *
agaatcATGTTCTTCTTCTTCtag
M F F F F *
agaatcATGTTCTTCTTCTTCTTCTTCtag
M F F F F F F *
agaatcATGTTCTTCTTCTTCTTCTTCTTCTTCtag
M F F F F F F F F *
agaatcATGTTCTTCTTCTTCTTCTTCTTCTTCTTCTTCtag
M F F F F F F F F F F *

**Figure S11**

**Effect of uORF length truncations on homopolymeric glutamic acid, arginine, and phenylalanine artificial uORFs.**

HEK293T cells were transfected with various pDLR *HOXA11* 5′ UTR constructs and luciferase activity determined 24 hours post transfection. *HOXA11* WT and a uORF-disrupted (TTG) mutant were included as controls. Sequential truncations (i.e. with 1, 2, 4, 6, 8, and 10 amino acid residues following the initial methionine) of homoploymeric artificial uORFs were assessed for (**A**) glutamic acid, (**B**) arginine, and (**C**) phenylalanine. Values are mean+SEM ($n$=4), and were scaled such that the mean of the WT control group was returned to a value of 1. Statistical significance was determined by one-way ANOVA with Bonferroni *post hoc* test. **$P<0.01$, ***$P<0.001$.

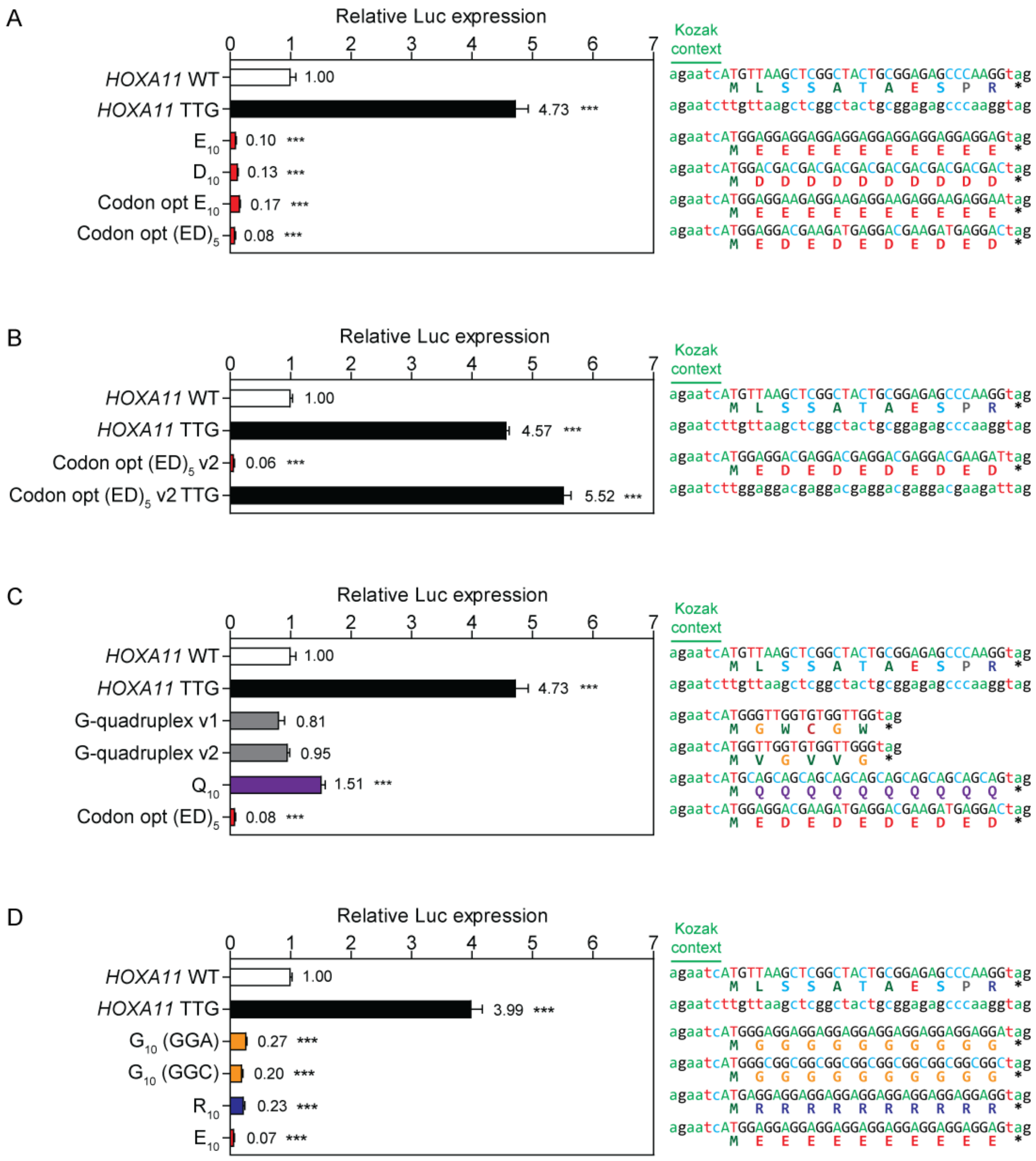
A
Relative Luc expression
0 1 2 3 4 5 6 7
HOXA11 WT 1.00
HOXA11 TTG 4.73 ***
E10 0.10 ***
D10 0.13 ***
Codon opt E10 0.17 ***
Codon opt (ED)5 0.08 ***
Kozak context
agaatcATGTTAAGCTCGGCTACTGCGGAGAGCCCAAGGtag
M L S S A T A E S P R *
agaatcttgttaagctcggctactgcggagagcccaaggtag
agaatcATGGAGGAGGAGGAGGAGGAGGAGGAGGAGGAGGtag
M E E E E E E E E E E *
agaatcATGGACGACGACGACGACGACGACGACGACGACtag
M D D D D D D D D D D *
agaatcATGGAGGAAGAGGAAGAGGAAGAGGAAGAGGAAtag
M E E E E E E E E E E *
agaatcATGGAGGACGAAGATGAGGACGAAGATGAGGACtag
M E D E D E D E D E D *
B
Relative Luc expression
0 1 2 3 4 5 6 7
HOXA11 WT 1.00
HOXA11 TTG 4.57 ***
Codon opt (ED)5 v2 0.06 ***
Codon opt (ED)5 v2 TTG 5.52 ***
Kozak context
agaatcATGTTAAGCTCGGCTACTGCGGAGAGCCCAAGGtag
M L S S A T A E S P R *
agaatcttgttaagctcggctactgcggagagcccaaggtag
agaatcATGGAGGACGAGGACGAGGACGAGGACGAAGATtag
M E D E D E D E D E D *
agaatcttggaggacgaggacgaggacgaggacgaagattag
C
Relative Luc expression
0 1 2 3 4 5 6 7
HOXA11 WT 1.00
HOXA11 TTG 4.73 ***
G-quadruplex v1 0.81
G-quadruplex v2 0.95
Q10 1.51 ***
Codon opt (ED)5 0.08 ***
Kozak context
agaatcATGTTAAGCTCGGCTACTGCGGAGAGCCCAAGGtag
M L S S A T A E S P R *
agaatcttgttaagctcggctactgcggagagcccaaggtag
agaatcATGGGTTGGTGTGGTTGGtag
M G W C G W *
agaatcATGGTTGGTGTGGTTGGGtag
M V G V V G *
agaatcATGCAGCAGCAGCAGCAGCAGCAGCAGCAGCAGtag
M Q Q Q Q Q Q Q Q Q Q *
agaatcATGGAGGACGAAGATGAGGACGAAGATGAGGACtag
M E D E D E D E D E D *
D
Relative Luc expression
0 1 2 3 4 5 6 7
HOXA11 WT 1.00
HOXA11 TTG 3.99 ***
G10 (GGA) 0.27 ***
G10 (GGC) 0.20 ***
R10 0.23 ***
E10 0.07 ***
Kozak context
agaatcATGTTAAGCTCGGCTACTGCGGAGAGCCCAAGGtag
M L S S A T A E S P R *
agaatcttgttaagctcggctactgcggagagcccaaggtag
agaatcATGGGAGGAGGAGGAGGAGGAGGAGGAGGAGGAtag
M G G G G G G G G G G *
agaatcATGGGCGGCGGCGGCGGCGGCGGCGGCGGCGGCtag
M G G G G G G G G G G *
agaatcATGAGGAGGAGGAGGAGGAGGAGGAGGAGGAGGtag
M R R R R R R R R R R *
agaatcATGGAGGAGGAGGAGGAGGAGGAGGAGGAGGAGGtag
M E E E E E E E E E E *

**Figure S12**

**Effect of codon optimization, RNA structure, and GC content on poly-acid artificial uORF activity.** HEK293T cells were transfected with various pDLR *HOXA11* 5′ UTR constructs and luciferase activity determined 24 hours post transfection. *HOXA11* WT and a uORF-disrupted (TTG) mutant were included as controls. The following mutants were tested: (**A**) A codon optimized version of the highly-repressive $E_{10}$ whereby sequence complexity was increased, and alternating glutamic and aspartic acids, $(ED)_5$ which was also codon-optimized to maximize sequence complexity. D10 was included as an additional control. (**B**) The highly repressive activity of a second codon-optimized (to remove an internal ATG trinucleotide) $(ED)_5$ mutant was tested by mutating its start codon to TTG. (**C**) Artificial uORFs containing potential for RNA secondary structure formation: a G-quadruplex sequence in two reading frames, and poly-glutamine ($Q_{10}$) where compared against the codon-optimized $(ED)_5$. (**D**) To test the effect of GC content, two variants of poly-glycine were tested; i.e. $(GGA)_{10}$ and $(GGC)_{10}$, relative to $E_{10}$ or $R_{10}$ controls. Values are mean+SEM (*n*=4), and were scaled such that the mean of the WT control group was returned to a value of 1. Statistical significance was determined by one-way ANOVA with Bonferroni *post hoc* test, ***$P$<0.001.

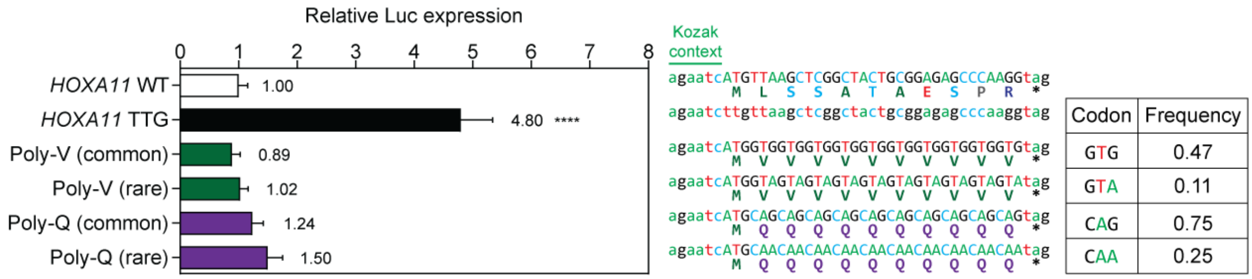


**Figure S13**

**uORF repressive activity is unaffected by rare codon usage.**

HEK293T cells were transfected with various pDLR *HOXA11* 5′ UTR constructs and luciferase activity determined 24 hours post transfection. *HOXA11* WT and a uORF-disrupted (TTG) mutant were included as controls. Homopolymeric artificial uORFs were tested consisting of valine or glutamine, using either rare or common codons. The frequency of codon usage for each codon is indicated. Values are mean+SEM (*n*=4), and were scaled such that the mean of the WT control group was returned to a value of 1. Statistical significance was determined by one-way ANOVA with Bonferroni *post hoc* test. ****$P$<0.0001.

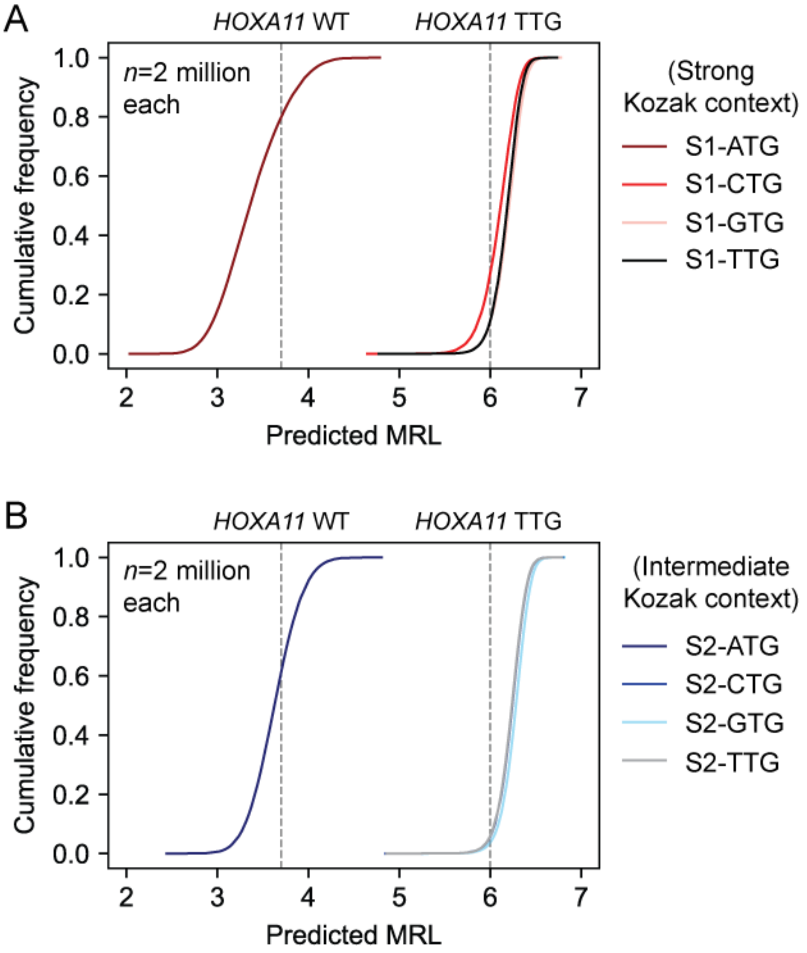


C

| *D*-statistic | S1-ATG | S1-CTG | S1-GTG | S1-TTG | S2-ATG | S2-CTG | S2-GTG | S2-TTG |
|---|---|---|---|---|---|---|---|---|
| S1-ATG | - | 1.000 | 1.000 | 1.000 | 0.362 | 1.000 | 1.000 | 1.000 |
| S1-CTG | 1.000 | - | 0.223 | 0.192 | 1.000 | 0.326 | 0.419 | 0.317 |
| S1-GTG | 1.000 | 0.223 | - | 0.062 | 1.000 | 0.105 | 0.214 | 0.095 |
| S1-TTG | 1.000 | 0.192 | 0.062 | - | 1.000 | 0.158 | 0.274 | 0.146 |
| S2-ATG | 0.362 | 1.000 | 1.000 | 1.000 | - | 1.000 | 1.000 | 1.000 |
| S2-CTG | 1.000 | 0.326 | 0.105 | 0.158 | 1.000 | - | 0.123 | 0.012 |
| S2-GTG | 1.000 | 0.419 | 0.214 | 0.274 | 1.000 | 0.123 | - | 0.134 |
| S2-TTG | 1.000 | 0.317 | 0.095 | 0.146 | 1.000 | 0.012 | 0.134 | - |

$P < 6.02 \times 10^{-117}$ for all pairwise comparisons

**Figure S14**

**Analysis of near-cognate uORF start codons using a machine learning approach.**

The S1 and S2 simulated 5′ UTR sets (**Figure 6A**) were further modified to investigate the effect of non-ATG, near-cognate start codons on uORF activity. Each simulation in S1 (Strong Kozak context) and S2 (intermediate Kozak context) set contains a single uORF (starting with an ATG) with randomized sequences embedded within the *HOXA11* 5′ UTR (i.e. S1-ATG and S2-ATG). Variation simulation sets were generated in which the ATG start codon was substituted for CTG, GTG, or TTG to gene for both S1 and S2 (each set contains 2 million simulations). Predicted mean ribosome load (MRL) values were generated and visualized by CDF plots for (**A**) S1-based simulation sets, and (**B**) S2-based simulation sets. Predicted MRL values for *HOXA11* WT and a uORF-disrupted (TTG) simulations are indicated with vertical dotted lines. (**C**) Table of pairwise Kolmogorov-Smirnov *D*-statistics for all comparisons.

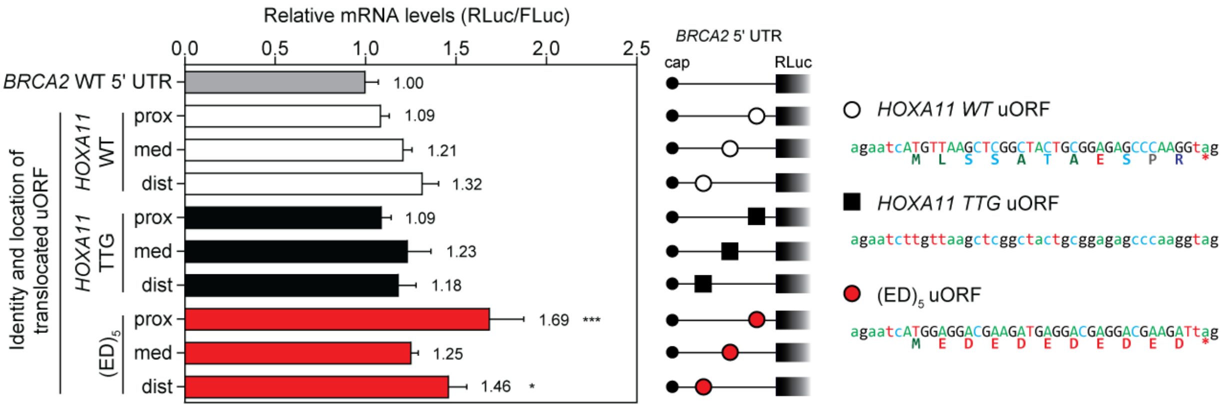


**Figure S15**

**Effect of uORF position on transcript levels.**

HEK293T cells were transfected with pDLR constructs as described in **Figure 8** and relative luciferase transcript levels were determined by RT-qPCR. RLuc expression was normalized to Fluc transcript levels and relative luciferase transcript levels determined 24 hours post transfection. Key mutated bases are highlighted in yellow. Values are mean+SEM, (*n*=3), and were scaled such that the mean of the WT control group was returned to a value of 1. Statistical significance was assessed by one-way ANOVA and Bonferroni *post hoc* test. *$P$<0.05, ***$P$<0.001.

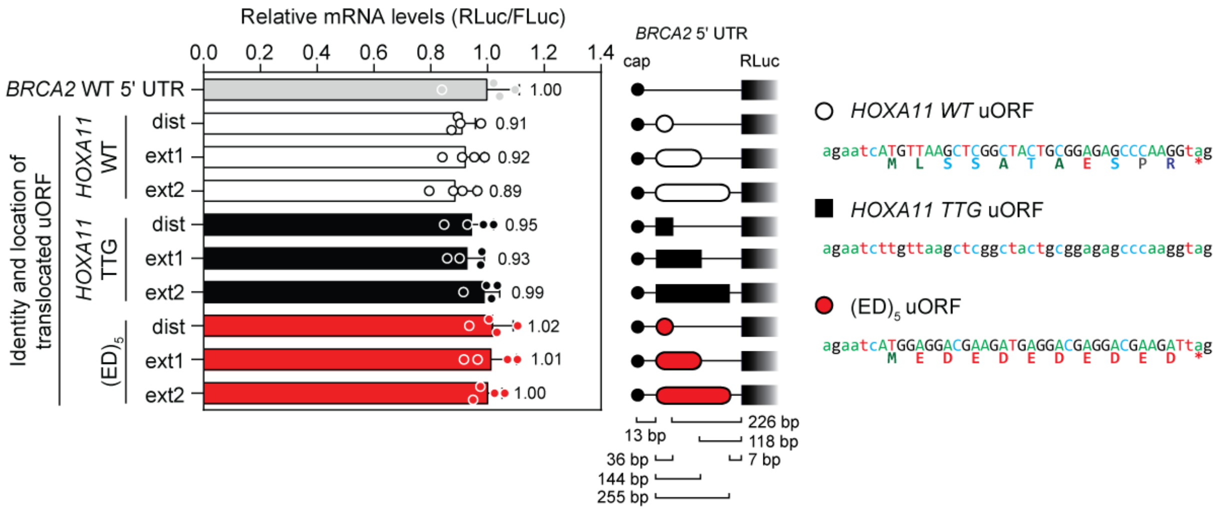


**Figure S16**

**Effect of uORF stop position on transcript levels.**

HEK293T cells were transfected with pDLR constructs as described in **Figure 8** and relative luciferase transcript levels were determined by RT-qPCR. RLuc expression was normalized to Fluc transcript levels and relative luciferase transcript levels determined 24 hours post transfection. Key mutated bases are highlighted in yellow. Values are mean+SD, ($n$=4 independent experiments), and were scaled such that the mean of the WT control group was returned to a value of 1. Statistical significance was assessed by one-way ANOVA and Bonferroni *post hoc* test.

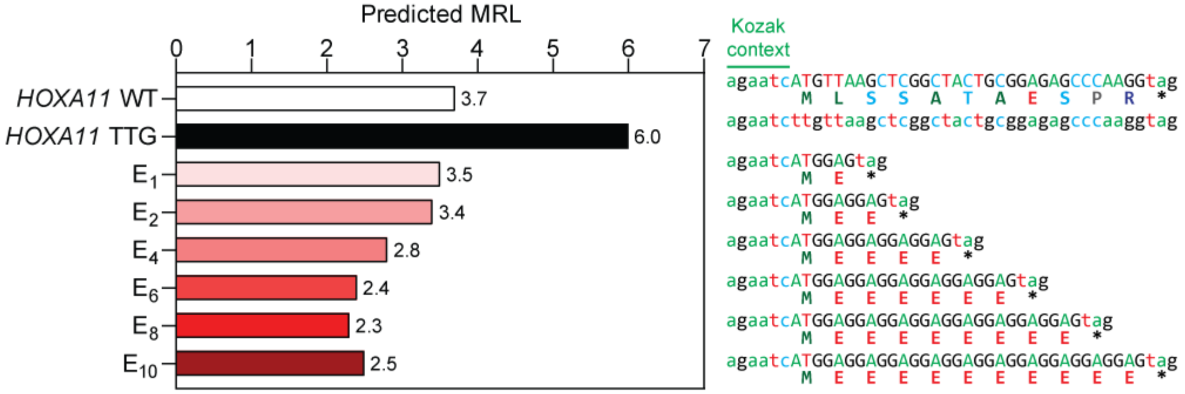


**Figure S17**

**Mean ribosome load predictions for homopolymeric glutamic acid artificial uORFs.**

Optimus 5-Prime was used to predict mean ribosome load (MRL) values for homopolymeric glutamic acid (Poly-E) artificial uORFs with sequential truncations (i.e. with 1, 2, 4, 6, 8, and 10 amino acid residues following the initial methionine), for which there is dual luciferase reporter experimental data (**Figure S11**). Predicted MRL values for *HOXA11* WT and a uORF-disrupted (TTG) simulations were included as controls.
